# Bayesian multilevel hidden Markov models identify stable state dynamics in longitudinal recordings from macaque primary motor cortex

**DOI:** 10.1101/2022.10.17.512024

**Authors:** Sebastien Kirchherr, Sebastian Mildiner Moraga, Gino Coudé, Marco Bimbi, Pier F Ferrari, Emmeke Aarts, James J Bonaiuto

## Abstract

Neural populations, rather than single neurons, may be the fundamental unit of cortical computation. Analyzing chronically recorded neural population activity is challenging not only because of the high dimensionality of activity in many neurons, but also because of changes in the recorded signal that may or may not be due to neural plasticity. Hidden Markov models (HMMs) are a promising technique for analyzing such data in terms of discrete, latent states, but previous approaches have either not considered the statistical properties of neural spiking data, have not been adaptable to longitudinal data, or have not modeled condition specific differences. We present a multilevel Bayesian HMM which addresses these shortcomings by incorporating multivariate Poisson log-normal emission probability distributions, multilevel parameter estimation, and trial-specific condition covariates. We applied this framework to multi-unit neural spiking data recorded using chronically implanted multi-electrode arrays from macaque primary motor cortex during a cued reaching, grasping, and placing task. We show that the model identifies latent neural population states which are tightly linked to behavioral events, despite the model being trained without any information about event timing. We show that these events represent specific spatiotemporal patterns of neural population activity and that their relationship to behavior is consistent over days of recording. The utility and stability of this approach is demonstrated using a previously learned task, but this multilevel Bayesian HMM framework would be especially suited for future studies of long-term plasticity in neural populations.

## 1. Introduction

The accelerating scale of recorded neurophysiological data has been accompanied by a shift from the view that individual neurons are the basic unit of computation, to one that explains computational processes in terms of the dynamics of neural populations (Saxena & Cunningham, 2019). Electrophysiology studies routinely record data from ever larger neural populations using chronically implanted multi-electrode arrays. These datasets are challenging to analyze because of their dimensionality, but recent evidence suggests that neural population activity in low dimensional subspaces, or manifolds, may be more important in neural computation than the firing of particular single neurons (Barack & Krakauer, 2021; Gallego et al., 2017, 2018; Humphries, 2021; Saxena & Cunningham, 2019). Chronically implanted electrode arrays present an additional analysis hurdle because the recorded signals can change dramatically over weeks and months of recording because of electrode drift or degradation (Barrese et al., 2016), or task-induced plasticity (Dayan & Cohen, 2011). The ability to understand long term learning mechanisms requires such longitudinal recording, but analysis of the resulting data at the individual neuron level is hindered by the difficulty in tracking single neurons over such long periods.

Hidden Markov Models (HMMs; Rabiner, 1989; Zucchini et al., 2017) are a particularly promising approach for analyzing the activity of large neural populations (e.g., Gat et al., 1997; Kemere et al., 2008; Mazurek et al., 2018; Ponce-Alvarez et al., 2012; Radons et al., 1994; Sadacca et al., 2016; Seidemann et al., 1996; van Kempen et al., 2021; Xydas et al., 2011). HMMs are probabilistic models which infer unobservable (hidden) “states” from a temporal sequence of observable data. These models can therefore be used to infer discretized lower dimensional state spaces that represent high dimensional neural activity. Given a suitable observational model governing the emission of observable data from unobservable states, HMMs do not require trial averaging or smoothing, making them suitable for trial-by-trial analyses.

Movements unfold in a specific spatiotemporal sequence, making it difficult to disentangle the underlying cognitive and motor mechanisms of goal-directed movements in which planning and feedforward and feedback-based control are fundamentally intertwined. The primary motor cortex is the largest source of corticospinal projections, making it uniquely situated to integrate these mechanisms and translate them to muscle coordination of the arm and hand. Because HMMs can identify discrete changes in continuous multivariate time series (Cunningham & Yu, 2014), they are a promising technique for decomposing dynamic M1 population activity into discrete states corresponding to the underlying cognitive and motor mechanisms of goal-directed movement. Previous HMM-based approaches to analyzing neural spiking activity have shown that such models can identify similar patterns of neural population activity in premotor cortex between execution and observation of the same actions (Mazurek et al., 2018), accurately predict changes in movement velocity in the primary motor cortex (Kadmon Harpaz et al., 2019), and segment baseline, plan, and peri-movement epochs from dorsal premotor cortex activity (Kemere et al., 2008). However, these approaches typically use statistical models with assumptions that are unmet by neural spiking activity, and in general have not considered non-stationary patterns of activity over long term recordings.

In the current study, we present a novel Bayesian multilevel HMM (e.g., de Haan-Rietdijk et al., 2017; Raffa & Dubin, 2015; Rueda et al., 2013; K. Shirley et al., 2016; Zhang & Berhane, 2014) adapted to longitudinal spiking data recorded from multi-electrode arrays. To accommodate the spike counts in multiple electrodes along with temporal overdispersion in the activity patterns, a multivariate Poisson log-normal emission distribution is adopted, with individual, trial-based covariates on the transition distribution. Each individual trial is thus allowed to have a specific sequence of hidden states, along with a specific set of transition probabilities and emission parameters via parametric random effects (i.e., trial-specific means) following a lognormal distribution centered on the group-level means.

We trained the model on spiking neural data recorded from the primary motor cortex of the macaque monkey during a reaching, grasping, and placing task. State transitions were tightly linked to task behavioral events, and state dynamics were closely aligned with the duration of intervals between events. We show that the states identified by the HMM represent fast transitions between distinct spatiotemporal patterns of neural population activity which are generated during specific phases of the reach, grasp, and place task, and likely correspond to different motor control processes.

## 2. Materials and methods

### 2.1. Multilevel Bayesian hidden Markov model

HMMs identify latent states from observable data by assuming that each state is only dependent on its previous state (the Markov assumption), and that only one state actively generates observations (emissions) at any given time point (i.e., states are mutually exclusive). The fitted model parameters include transition probabilities, which define the likelihood of transitioning from any given state to any other state, and emission probabilities, which govern the generation of observable data within each state. Classic frequentist HMMs applied to neural data have used a univariate multinomial distribution for emission probabilities (Bollimunta et al., 2012; Mazurek et al., 2018; Diomedi et al., 2021), encoding which of *N* neurons is spiking in each time bin, and choosing a random neuron if multiple neurons spike in the same bin. Such an approach does not capture the multivariate nature of neural population activity measured with high density recordings, nor the statistical distribution of neural spiking, and breaks down as larger neural populations are recorded from. Previous HMM frameworks which have used multivariate and/or Poisson emission probabilities (Radon, 1994; Siedemann, 1996; Kemere, 2008; Xydas, 2011; Ponce-Alvarez, 2012), have been single-level, and thus unable to model day-to-day variability in longitudinal neural recordings. We therefore used a multilevel Bayesian hidden Markov model framework (K. E. Shirley et al., 2012) to mitigate these shortcomings, modeling trial-specific random effects in transition and emission probabilities, multivariate Poisson log-normal emission probability distributions, and trial-specific condition covariates.

The adoption of the multilevel framework therefore has a number of advantages over previous non-hierarchical approaches. First, all the information available in the full data set is used for the estimation of each trial-specific parameter, such that parameter estimation on trials consisting of a lower number of observations benefits from trials with more observations (Hox et al., 2018; Schoot & Miočević, 2020). As a result, the hierarchical structure of the model produces a regularizing effect on the estimation of trial-specific parameters, making them more robust to outliers (Gelman & Pardoe, 2006). Second, the inclusion of parametric random effects as part of the hierarchical structure of the model not only offers a vehicle for the allocation of variability between trials, but also avoids the statistical caveats of ignoring the nested structure of the electrophysiological recordings such as the underestimation of standard errors (Aarts et al., 2014; Hox et al., 2018; Schoot & Miočević, 2020). Finally, the multilevel framework provides each trial with individual parameters including an individual trajectory over time, while ensuring that the meaning of each state remains the same across trials. This between-trial state consistency avoids the need for post-hoc state matching between trials, as would be necessary if separate single-level models were fit to data from each day or recording session.

In addition, the Bayesian HMM framework has several advantages over previous implementations relying on maximal likelihood estimation and expectation-maximization paradigms. First, it allows for the inclusion of trial-specific random effects in both transition and emission probabilities, which would rapidly become computationally intractable in alternative frameworks as the number of random effects (trials) and model complexity grow (Altman, 2007; de Haan-Rietdijk et al., 2017). Second, the Bayesian estimation procedure produces useful byproducts, at no extra computational cost, such as the local decoding of the states (the most likely state at each point in time) and credibility intervals (Bayesian analogue to confidence intervals) on all model parameters, avoiding the necessity for additional computationally intensive steps for estimation of parameter standard errors. Finally, although non-informative priors are most commonly used, adopting (weakly) informative priors allows the possibility of updating the model with new data as it becomes available.

#### 2.1.1 Model specification of a basic HMM

The likelihood *L_T_* of a basic hidden Markov model (HMM) consisting of a single time series of observations *o*_1:*T*_ and a first-order Markov chain of *m* hidden states *s*_1:*T*_ can be defined in terms of three basic components

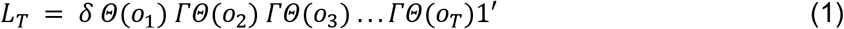

where *δ* is a vector of dimensions (1 × *M*) with the initial probabilities of the states *s* ∈ {*S*_1_,…,*S_m_*}, *Γ* is a transition probability matrix of dimensions (*M* × *M*) denoting the transition probabilities *γ_ijt_* = *P*(*s_t_* = *S_j_* | *s*_*t*-1_ = *S_i_*) which is assumed to be homogeneous over the occasions *t, Θ* is the emission distribution consisting of a diagonal matrix of dimensions (*M* × *M*) with the *i*th diagonal element denoting the emission probability density *θ_i_* = *P*(*o_t_* = *O_t_* | *s_t_* = *S_i_*) which can take any arbitrary form, and 1’ is a column vector of *M* elements of value one. In the context of this study, a Poisson emission distribution is assumed for the observed spike counts *o_t_*, given the hidden state *s_t_,* such that *P*(*o_t_* = *O_t_* | *s_t_* = *S_i_*) ≡ *Poisson*(*λ_i_*). *λ* is the diagonal matrix of dimensions (*M* × *M*) with its diagonal elements denoting the Poisson parameters (expected spike counts) for the *M* hidden states *S*.

Although the model has been defined for a single observed sequence, it can readily be extended to multivariate data with the convenient assumption that the multiple sequences (e.g., observations from multi-electrode probes) are conditionally independent given the hidden states (e.g., Zucchini et al., 2017). That is, we specify the joint distribution *P*(*o*_1*t*_ = *O*_1*t*,*o*_2*t*__ = *O*_2*t*_,…,*o_kt_* = *O_kt_*) for the *k* ∈ {1,2,…,*K*} state-dependent emission distributions as the product of the *K* marginal state-dependent emission probability densities *P*(*o_kt_* = *O_kt_* | *s_t_* = *S_i_*):

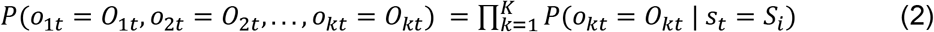

As a result, the likelihood of the general multivariate HMM takes the form:

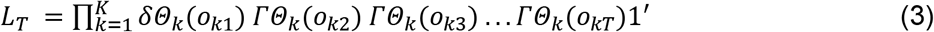

A basic HMM of this specification can be easily estimated using direct likelihood maximization (e.g., Zucchini et al., 2017), following a computationally efficient recursive implementation of the Expectation-Maximization algorithm (Dempster et al., 1977) known as the Baum-Welch algorithm (Baum et al., 1970; Zucchini et al., 2017), or via the Bayesian estimation framework as we do here (e.g., Scott, 2002).

#### 2.1.2 Model specification of the Bayesian multilevel HMM

In the current implementation of the multilevel HMM, *γ_nij_*, the probability for the individual *n* ∈ {1,..,*N*} (normally an individual, but here each trial) of switching between a state *i* and state *j*, at occasion *t*, follows a Multinomial Logit model

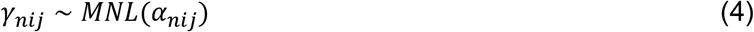

Hence, each trials’ probability to transition from hidden state *i* ∈ {1,..,*M*} to state *j* ∈ {1,..,*M*} is modeled using *M* batches of *M* – 1 random intercepts, *α_nij_*. That is,

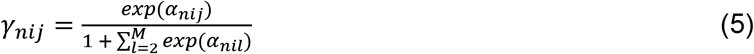

Where the numerator is set equal to *1* for *j* = 1, making the first state of every row of the transition probability matrix *Γ* the baseline category.

For the Bayesian implementation we adopt a group-level state-specific multivariate Normal prior distribution on the trial-specific parameters *α_nij_*, with mean vector 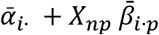 of (*M* – 1) elements, and covariance –denoting the covariance between the *M* – 1 state *i*-specific intercepts over trials. Thus, in addition to the group-level intercepts 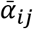, we have *M* matrices of *p* * (*M* – 1) fixed regression coefficients, 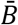, where *p* denotes the number of used covariates. The columns of 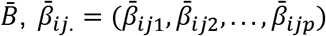, are used to model the random intercepts denoting the probability of transitioning from state *i* to state *j* given *p* individual-level covariates.

A convenient hyper-prior on the parameters of the group-level prior distribution is a multivariate Normal distribution for the mean vector 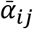 and the fixed regression coefficients 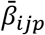, and an Inverse Wishart distribution for the covariance *Ψ_ij_*. That is,

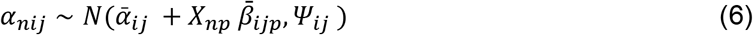

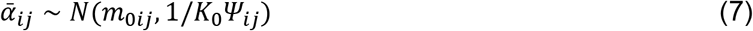

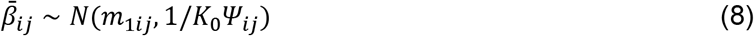

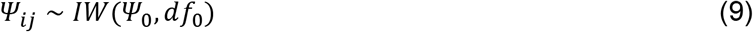

The parameters *m*_0*ij*_, *m*_1*ij*_, and *K*_0_ denote the values of the parameters of the hyper-prior on the group (mean) vector 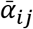, and 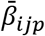, respectively. Here, *m*_0*ij*_ and *m*_1*ij*_ represent vectors of means, and *K*_0_ denotes the number of observations (i.e., the number of hypothetical prior trials) on which *m*_0*ij*_ and *m*_1*ij*_ are based. The parameters *Ψ*_0_ and *df*_0_, respectively, denote values of the covariance and the degrees of freedom of the hyper-prior for the Inverse Wishart distribution on the group variance *Ψ_ij_* of the trial-specific random intercepts *α_nij_*.

Just as for the basic HMM introduced in section 2.1.1, spike count data *o_nkt_* follow Poisson emission distributions, but here with trial-specific parameters *λ_nik_* conditionally independent on the state *i* ∈ {1,.., *M*}. That is,

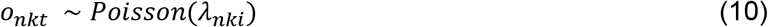

for electrode (the dependent variable) *k* ∈{1,..,*K*}, on time point *t*∈{1,..,*T*}, and trial *n* ∈ {1,.., *N*}, given the state *i*.

We adopt a group-level state-specific log-Normal prior distribution on the trial-specific parameters *λ_nki_*, with mean vector 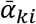 of *M* elements, and variance *τ_ki_*. Note that for the emission distribution no covariates have been used to model the variability between the trial-specific, electrode-specific, state-specific parameters *λ_nki_*. The complete specification of the emission distribution adopted is detailed in probabilistic notation below, along with conveniently defined hyper-priors on the parameters of the model

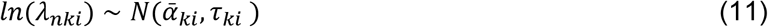

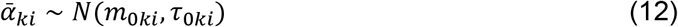

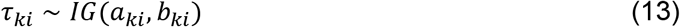

The parameters *m*_0*ki*_ denote the *a priori* expectations on the group-level mean counts on the logarithmic scale, along with the *a priori* expectations on their variability, *τ*_0*ki*_. Finally, *α_ki_, b_ki_* are the hyper parameters denoting the shape and rate of the hyper prior on the between-individual variance *τ_ki_*.

Note that using a Normal prior on the log-transformation of the trial-specific, electrode-specific, state-specific Poisson parameters *λ_nki_* is equivalent to specifying a log-Normal prior on the untransformed Poisson parameter. In addition, as previously mentioned, because *λ_nki_* are assumed to be conditionally independent given the state, no covariance matrix is estimated for the covariance between dependent variables.

### 2.2 Reach-to-grasp task

Two adult female rhesus macaques (*Macaca mulatta*) trained on a reach and grasp task served as the subjects. The animal handling as well as surgical and experimental procedures complied with European guideline (2010/63/UE) and authorized by the French Ministry for Higher Education and Research (project # 2016112713202878) in force on the care and use of laboratory animals, and were approved by the ethics committee CELYNE (comité d’éthique Lyonnais pour les neurosciences expérimentale, C2EA 42). After initial training, we performed a sterile surgery to implant six floating multielectrode arrays (FMA, Microprobes for Life Science, Gaithersburg, MD, USA) in the right (monkey 1) or left (monkey 2) cortical hemisphere. Each array was comprised of 32 platinum/iridium electrodes (impedance 0.5 MΩ at 1 kHz) with lengths ranging from 1 to 6 mm, and with an inter-electrode spacing of 400 μm. One electrode array was implanted in the primary motor cortex (M1), two were implanted in the ventral premotor cortex (F5), one in the dorsal premotor cortex (F2), and two in the prefrontal cortex (45a and 46/12r), as estimated according to a previous magnetic resonance imaging scan. For the purposes of this study, we analyzed data from the M1 array of each monkey.

Throughout the task, the monkey sat in front of a table containing a handle and a semicircular groove. A metallic cube, the target, was placed at 11 cm from the handle, in a slot within a metallic grasping platform which could be placed within the groove (Figure 1B). Contact with the handle, target object, and the bottom of the groove was recorded by a circuit which detected changes in resistance. The task was programmed and controlled by EventIDE software (OkazoLab Ltd). Trials started when the monkey grasped the handle. After a variable delay period (500 - 1000 ms), an auditory go signal (900 Hz tone) instructed the monkey to reach for the target object, grasp it, and place it into the groove beside the grasping platform. The monkey was then rewarded with a few drops of water. Data was recorded from 1 second before the handle grasp until the reward. The task was run in blocks of 10 trials per condition (with the grasping platform placed in the left, right, or center of the groove). The left and right target positions were each 25° from the center of the semicircular groove, relative to the handle. The orientation of the slot in the grasping platform depended on the condition. In the center condition the slot was oriented at 0° with respect to the animal, and in the left and right conditions the slot was oriented at 45° to left and right, respectively. The monkey completed 2-4 blocks of 10 trials per day by condition (monkey 1: left: *M* = 17.1, *SD* = 5.4, center: *M* = 16.7, *SD* = 5.1, right: *M* = 16.7, *SD* = 7.2; monkey 2: left: *M* = 16.0, *SD* = 4.1, center: *M* = 13.1, *SD* = 4.4, right: *M* = 15.2, *SD* = 3.9), for a total of 25-70 trials per day (monkey 1: *M* = 50.5, *SD* = 13.3; monkey 2: *M* = 44.3, *SD* = 11.9). Any trial in which the handle was released before the go signal or the grasping movement was not properly executed within 3 seconds was aborted and not rewarded. Each monkey used the hand contralateral to the implanted array to perform the task (monkey 1: left hand, monkey 2: right hand).

**Figure 1.**
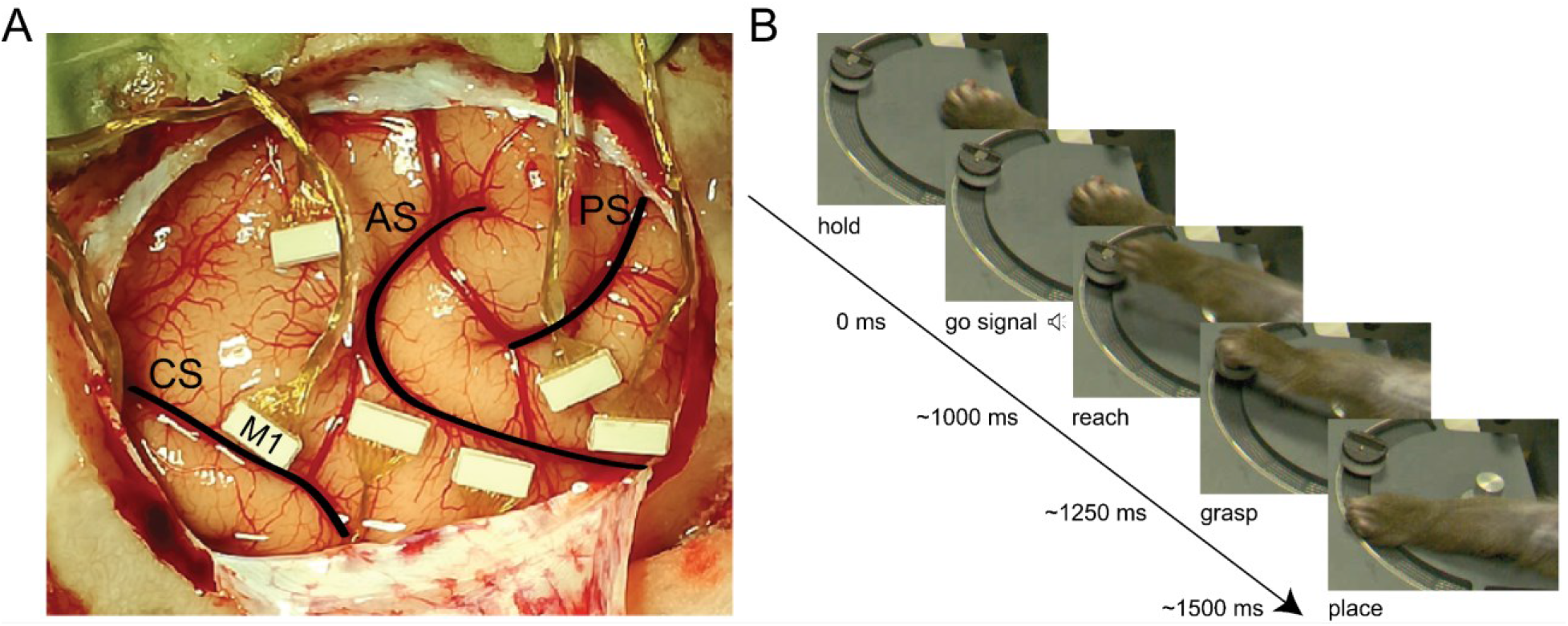
Experimental setup. **A)** Localization of the floating multi-electrode arrays (FMA) on the right cortical hemisphere of monkey 1. Six arrays were implanted in the frontal cortex, and here we analyze data from one which was located in the primary motor cortex (M1). Labels show the location of the arcuate sulcus (AS), central sulcus (CS), and principal sulcus (PS). **B)** The monkey performed a cued reach-to-grasp task. In each trial, the monkey was required to grasp a handle until the go signal, then reach and grasp a metallic cube with a precision grasp (using the index finger and thumb), and finally place it into a groove in the table.

### 2.3 Neural recordings

The wideband neural signal (bandpass filtered at 0.1 to 7500 kHz) was recorded at 30 kS/s, and amplified and digitized (16-bit; 0.192 μV resolution) with an Intan Tech-based (Intan Technologies, Los Angeles, CA, USA) open source acquisition system (Open Ephys; Siegle et al. 2017). This system uses a 256-channel Intan RHD2000 series acquisition board and 32-channel headstages (RHD2132). Spike detection was performed offline using Trisdesclous (Garcia & Pouzat,2015). The common reference was removed to reduce ambient noise. Spikes were then detected from each electrode using a threshold of 2 times the median absolute deviation (MAD), and analyzed as multi-unit activity (MUA) in 10 ms bins. All electrodes in which at least one well-isolated spike waveform was detected were selected for the following analyses. We thus used a sample of 21 electrodes out of 32 for monkey 1, and 25 out of 32 electrodes for monkey 2. Custom made detection panels were used to record the moments when the monkey’s hand released the handle, the hand contacted the target object, and when the object was placed in the groove. An Omniplex 16-channel recording system (Plexon, Dallas, TX, USA) was used to simultaneously record these behavioral events. Trials were discarded if the response time (time between the go signal and handle release) was less than 100 or greater than 1500 ms, the reach duration (time between handle release and object contact) was less than 100 or greater than 1000 ms, or the placing duration (time between object contact and placing the object in the groove) was less than 100 or greater than 1200 ms, leaving 19 - 68 trials per day for monkey 1 (*M* = 43.9, *SD* = 15.46, *N* = 439; left: *M* = 14.8, *SD* = 5.74; center: *M* = 14.4, *SD* = 5.15; right: *M* = 14.7, *SD* = 7.73), and 23 - 49 per day for monkey 2 (*M* = 38.3, *SD* = 9.87, *N* = 383; left: *M* = 14.2, *SD* = 3.91; center: *M* = 10.8, *SD* = 3.55; right: *M* = 13.3, *SD* = 3.37).

### 2.4 Model training

Model training was performed using R (R Core Team, 2021) and the developer version of the package *mHMMbayes* (Aarts, 2019). In *mHMMbayes*, models are fitted using a hybrid Metropolis within Gibbs MCMC algorithm, which expands on classic HMM implementations by using Bayesian estimation as outlined in Scott (2002). Estimation results from a conventional expectation-maximization HMM trained on aggregated spike count data were used as starting values for the MCMC chain (depmixS4; Visser & Speekenbrink, 2010). The Bayesian multilevel HMM was fit to binned (10 ms bin width) spike counts from each electrode used with trial-specific intercepts, and covariates encoding each trial’s condition. The model was fit with 4000 iterations, with the first 2000 being discarded to dissipate the effect of starting conditions (burn-in). In fitting the model, agnostic non-informative hyper-priors were specified for all group-level parameters. Convergence of all sample-level parameters was checked by verifying that the multivariate potential scale reduction factor, 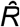, was lower than a threshold of 1.05 (Brooks & Gelman, 1998) for two additional chains with randomized starting values. The sequence of most likely states given the neural data, along with each state’s forward probabilities were determined for each trial with the Viterbi algorithm (Forney, 1973; Viterbi, 1967) based on trial-specific parameters.

Model selection was performed by comparing relative model fit using AIC (Akaike, 1974) and BIC (Schwarz, 1978). The ability of the selected model to reproduce the original multielectrode data was determined using Bayesian posterior predictive checks (PPCs; for more details see: Gelman & Shalizi, 2013; McElreath, 2020). For the PPCs, the fitted model was used to simulate 500 new data sets, after which we assessed the extent to which a set of summary statistics (mean, standard deviation, maximum counts, and proportion of zeros) on all electrodes (21 for monkey 1 and 25 for monkey 2) at the aggregated sample-level in the simulated data recovered the values empirically observed in the electrophysiological data.

The upper bound for the decoding accuracy of the neural states identified by the multilevel HMM was assessed with a small Monte Carlo simulation. The fitted model was used to simulate 100 new data sets with the same number of trials and observations per trial as the empirical data used to fit the model. The simulated data sets were then used to fit new multilevel HMMs, after which the accuracy of state decoding was evaluated.

### 2.5 Neural and behavioral data analysis

To compare state onset and offset times to behavioral event timings, we identified episodes lasting at least 100 ms in each trial where a particular state was identified as the most likely using the Viterbi algorithm. If a state was activated multiple times in a trial, we used the first activation. The partial Spearman’s correlation coefficient was used to relate state onset and offset times to each behavioral event time (movement onset, object contact, and placing), accounting for correlations between other event timings and the onset or offset of all other states. Statistical significance was tested using permutation tests (10,000 permutations) in which the event timings, but not the state onset and offset times, were shuffled in each iteration.

Response time (the time between the go and movement onset events), reach duration (time between the movement onset and object contact events), and placing duration (time between the object contact and placing events) were computed for each trial and compared between conditions using linear mixed models with condition (left, center, right) as a fixed effect and day-specific offsets as a random effect. The number of state activations per trial was given by the number of episodes where a state was identified as the most likely for at least 100 ms. To compute state lifetime, the first activation was used if a state was activated multiple times in a trial, as above. State activations and lifetime were compared across states and conditions using generalized linear mixed models (with Poisson distribution and log link function for activations, and Normal distribution and identity link function for lifetime) with condition, state, and their interaction as fixed effects, and trial nested within day offsets as random effects. The relationship between response time, reach duration, and placing duration and the number of activations and lifetime of each state was evaluated using partial Spearman correlation coefficients, accounting for all other behavioral durations and state activations or lifetimes.

Multi-unit firing rate activity around state transitions was compared by aligning firing rates to the onset and offset of the maximum duration activation of each state around a time window lasting from 100 ms prior to the onset to half of the mean state lifetime after onset, and from half the mean state lifetime prior to offset to 100 ms after offset. The firing rate from each electrode was then baseline-corrected using the mean firing rate in the 100 ms prior to the state onset. For each electrode, we compared the mean, baseline-corrected firing rate in the 100 ms prior to state onset to that in the first half of the state lifetime using linear mixed models with state, time period (baseline or first half of lifetime), and their interactions as fixed effects, and trial nested within day offsets as random effects. In the same way, we compared the mean, baseline-corrected firing rate in the first half of the state lifetime to that in the second half.

Temporal and spatial shuffling was performed by permuting the data either by time point or by electrode, 100 times. The resulting permuted data was run through the HMM to obtain forward probabilities and sequences of the most likely state. For each iteration, the partial Spearman coefficient between state onsets and offsets and the behavioral events (accounting for correlations between other event timings and the onset or offset of all other states, as above) was evaluated, and the resulting coefficients were used as the null distribution for comparison with the corresponding unshuffled coefficients.

All mixed models were run in R (R Core Team, 2021) using *lme4* (v1.1.26; Bates et al., 2018). Factor significance was determined using type II Wald X^2^ tests using the *car* package (v3.0.10; Fox et al., 2019), and pairwise Tukey-corrected follow-up tests were performed using estimated marginal means using the *emmeans* package (v1.5.3; Lenth et al., 2020) with Kenward-Roger approximated degrees of freedom. Code for all analyses is freely available (https://github.com/danclab/motor_mea_mbhmm).

## 3. Results

Multiple multilevel Bayesian HMMs with 3 to 8 states were fit to single-trial multi-unit spiking activity (Figure 2A) recorded from the primary motor cortex of two macaque monkeys during a reach-grasp task. Model comparison (AIC and BIC) was used to select the optimal model, yielding a model with 6 states for monkey 1, and 5 states for monkey 2 (Figure S1). All group-level parameters in the optimal models met the convergence criterion.

**Figure 2.**
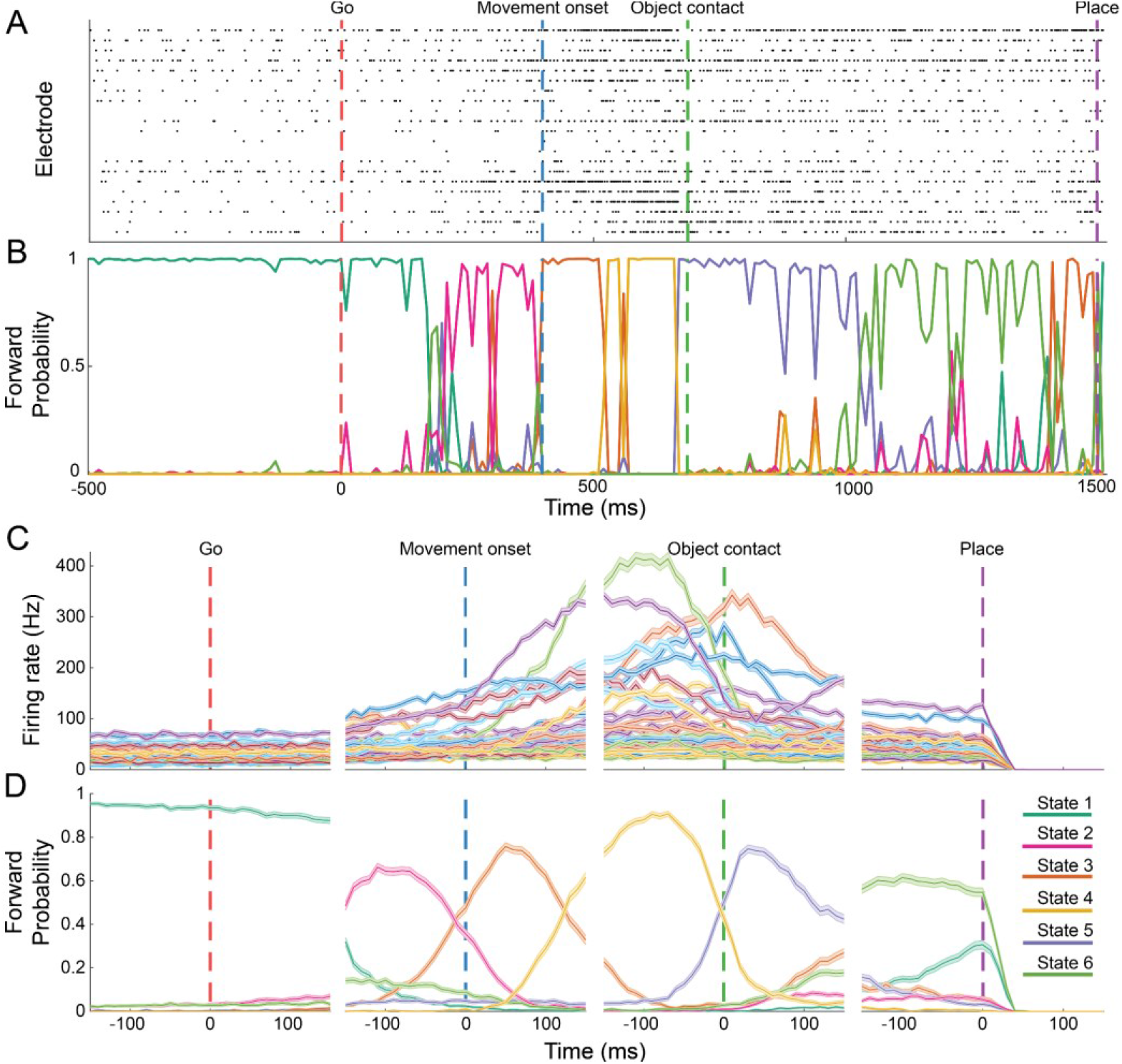
The HMM identifies states that correspond to different phases of the action. **A)** Raster plot of a single example trial representing the spiking activity from 21 electrodes from the M1 array of monkey 1. **B)** Forward probabilities for each state given by the model for the same trial shown in (A). **C)** Mean multi-unit firing rate of 21 electrodes from the M1 array of monkey 1 over 10 days (439 trials) aligned to behavioral events (go: go signal tone, hand movement onset: beginning of the reaching movement, object contact: grasping of the object, place: placing the object). **D)** Mean forward probabilities generated by the model for each state, averaged over the same trials and aligned to the same behavioral events as in (C).

### 3.1 Simulations

We started by evaluating the goodness of fit of the optimal models selected for the two monkeys with posterior predictive checks. The results of the posterior predictive checks (PPCs) show that the models were able to recover the overall patterns of electrical activity aggregated over trials for the 21 and 25 electrodes of the first (Figure S2) and second monkey (Figure S3), respectively, which indicates a good fit to the empirical data. In addition, for some of the electrodes, the PPCs reveal a small overestimation of the mean aggregated counts, coupled with a small underestimation of the proportion of bins with zero spike counts (e.g., monkey 1 electrodes 6, 10, and 21; Figure S2) which are consistent with a zero-inflated generative process. These minor deviations from good fit appear to be more pronounced for the data of the second monkey than for the first monkey.

Next, we explored the expected state decoding accuracy with a Monte Carlo simulation based on the modeling results for the optimal models on the two monkeys. The mean percent balanced accuracy averaged over trials and simulation repetitions of 91.6% (95%CI [91.3-92.0%]) for monkey 1 and 87.5% (95%CI [86.3-88.7%]) for monkey 2 (see Table S1 for extended performance metrics). The mean balanced accuracy varied across states, ranging from 73.0% for state 2 to 96.6% for state 1 in monkey 1 and from 60.8% for state 2 to 98.6% for state 3 in monkey 2 (Table S2). Given the good fit to the data (Figures S2 and S3), these results indicate that a high level of decoding accuracy can be expected for the empirical data.

### 3.2 Distinct M1 population activity states occur during different phases of the reach-to-grasp task

For each state, the model generates a forward probability, representing the probability at each point in time that the neural population is in that state given the MUA (Figure 2B). States were labeled in order of their trial-averaged order of occurrence. Trials had different durations due to variability in response time and movement kinematics, so in order to determine the task phase specificity of each state, we aligned both multi-unit firing rates and each state’s forward probability with each behavioral event (go signal, hand movement onset, object contact, and placing), and averaged over trials. The event-aligned multi-unit mean firing rate exhibited slow ramping dynamics prior to the movement onset, and peaking just before object contact (monkey 1: Figure 2C; monkey 2: Figure S4A). In contrast, the mean forward probabilities of each HMM state indicate that discrete state transitions tended to occur around the same time as the movement onset, object contact, and object placing behavioral events (Figure 2D). At the start of each trial, state 1 predominated, but sometime following the go signal, there was a transition to state 2, followed by a transition to state 3 after the hand movement onset. During the initial reach, there was a transition to state 4, and then to state 5 after the hand contacted the object. Finally, during the final reaching movement to place the object in the groove, there was a transition to state 6. This sequence of state transitions was similar for the 5 states identified from monkey 2 (Figure S4B). Importantly, behavioral event timings were not provided to the HMM during training, it was simply fit to the multi-unit spiking activity recorded during each trial. This suggests that the identified states represent actual states of the neural population corresponding to different processes involved in motor control, rather than trivially reflecting the temporal structure of the task.

Although it appears that state transitions occur in specific relation to behavioral events, this could have been a result of trial averaging. We therefore examined the temporal relationship between state transitions and behavioral events at the single trial level. In each trial, increases and decreases in state forward probabilities closely matched the timing of the movement onset, object contact, and placing events (monkey 1: Figure 3; monkey 2: Figure S4C). In each trial, there was a transition from state 1 to state 2 occurring before the hand movement onset, from state 2 to 3 closely locked to the hand movement onset, from 3 to 4 between the hand movement onset and object contact, from 4 to 5 locked to object contact, and from 5 to 6 between object contact and the placing event. We next used the Viterbi algorithm (Forney 1973; Viterbi 1967) to estimate the most likely state at each point in time, for every trial, from the forward probabilities. We used the sequence of most likely states to determine the onset and offset times of each state in every trial. We then ran a permutation test on the partial Spearman’s correlation coefficient (accounting for correlations between event timings and the onsets and offsets of other states) between the onset and offset time of each state and the timing of each behavioral event. This revealed that for monkey 1, the onset (*ρ* = 0.73, *p* < 0.001) and offset (*ρ* = 0.64, *p* < 0.001) of state 2 were most correlated with the movement onset time. Only the onset of state 3 (*ρ* = 0.46, *p* = 0.003), the offset of state 4 (*ρ* = 0.56, *p* < 0.001), and the onset of state 5 (*ρ* = 0.46, *p* = 0.005) were correlated with the object contact event. The offset of state 5 (*ρ* = 0.26, *p* = 0.037) and the onset of state 6 (*ρ* = 0.32, *p* = 0.005) were the transitions most correlated with the placing event. For monkey 2, the onset (*ρ* = 0.33, *p* < 0.001) and offset (*ρ* = 0.32, *p* < 0.001) of state 2 was were correlated with movement onset time, the onset of state 3 (*ρ* = 0.15, *p* = 0.042) and the onset (*ρ* = 0.24, *p* = 0.001) and offset (*ρ* = 0.30, *p* < 0.001) of state 4 were correlated with object contact, and the onset (*ρ* = 0.35, *p* < 0.001) and offset (*ρ* = 0.38, *p* < 0.001) of state 4 and the onset of state 5 (*ρ* = 0.15, *p* = 0.046) with the placing event. We then evaluated these correlations over the duration of the recordings, in groups of two days to ensure a large enough number of trials for each correlation. In both monkeys, these correlations were sustained over 10 days of recording (all *p* < 0.02; Figure 4). The onset and offset of various states was therefore tightly linked to the movement onset, object contact, and placing events at the single trial level, with earlier occurring states tending to be related to movement onset, states in the middle of the trial related to object contact, and states occurring later related to the placing event. Moreover, this linkage between states and behavioral events was stable over the entire period of recording.

**Figure 3.**
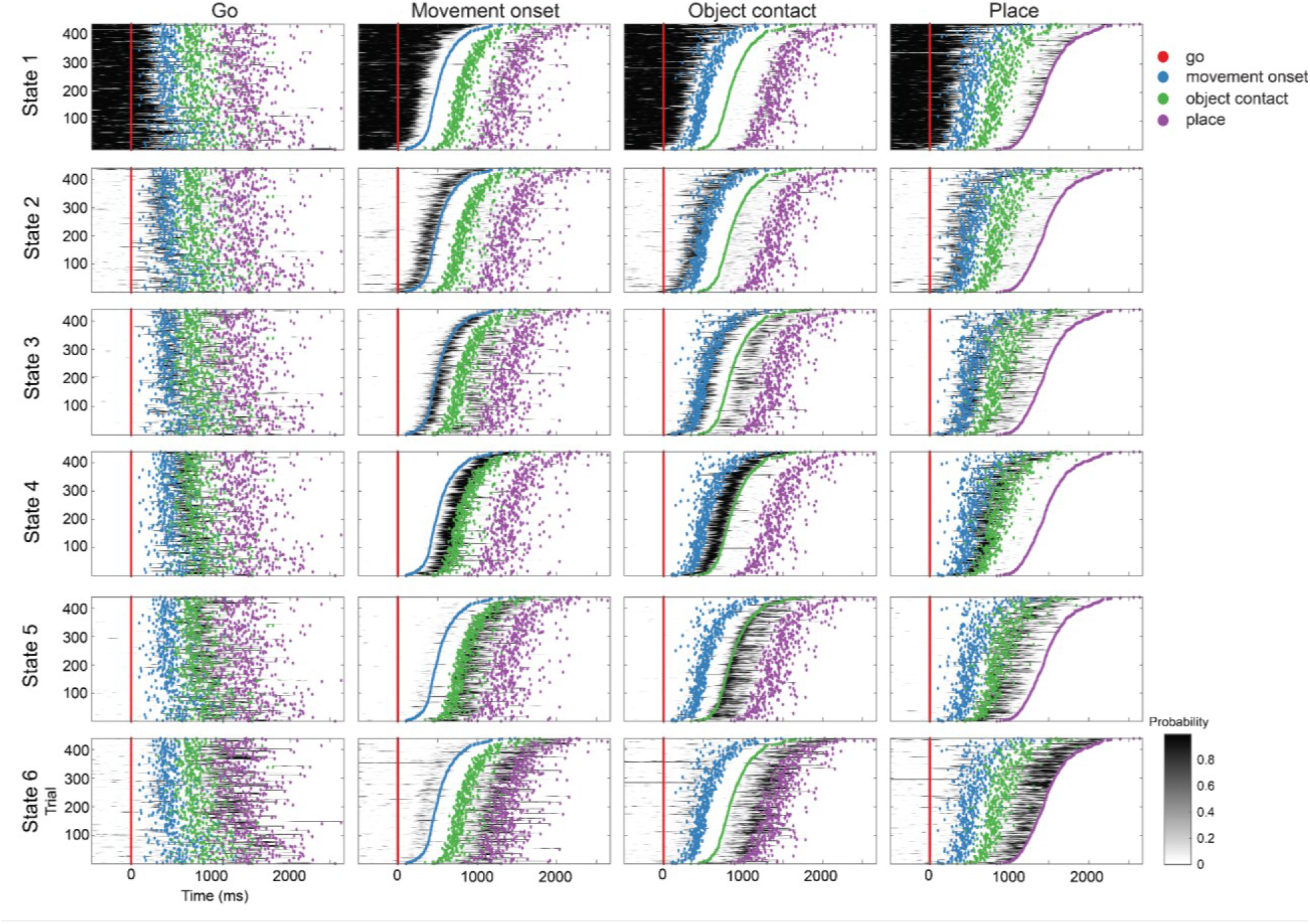
States are tightly linked to behavioral events at the single trial level. Each row shows the forward probabilities from one of the model states for each trial, over all days of recording, for monkey 1. Each column shows the trial forward probabilities sorted by one of the behavioral events (from left to right: go signal, hand movement onset, object contact, and placing). When trials are sorted by the appropriate event, the alignment between state forward probabilities and event timing can be clearly seen over all trials.

**Figure 4.**
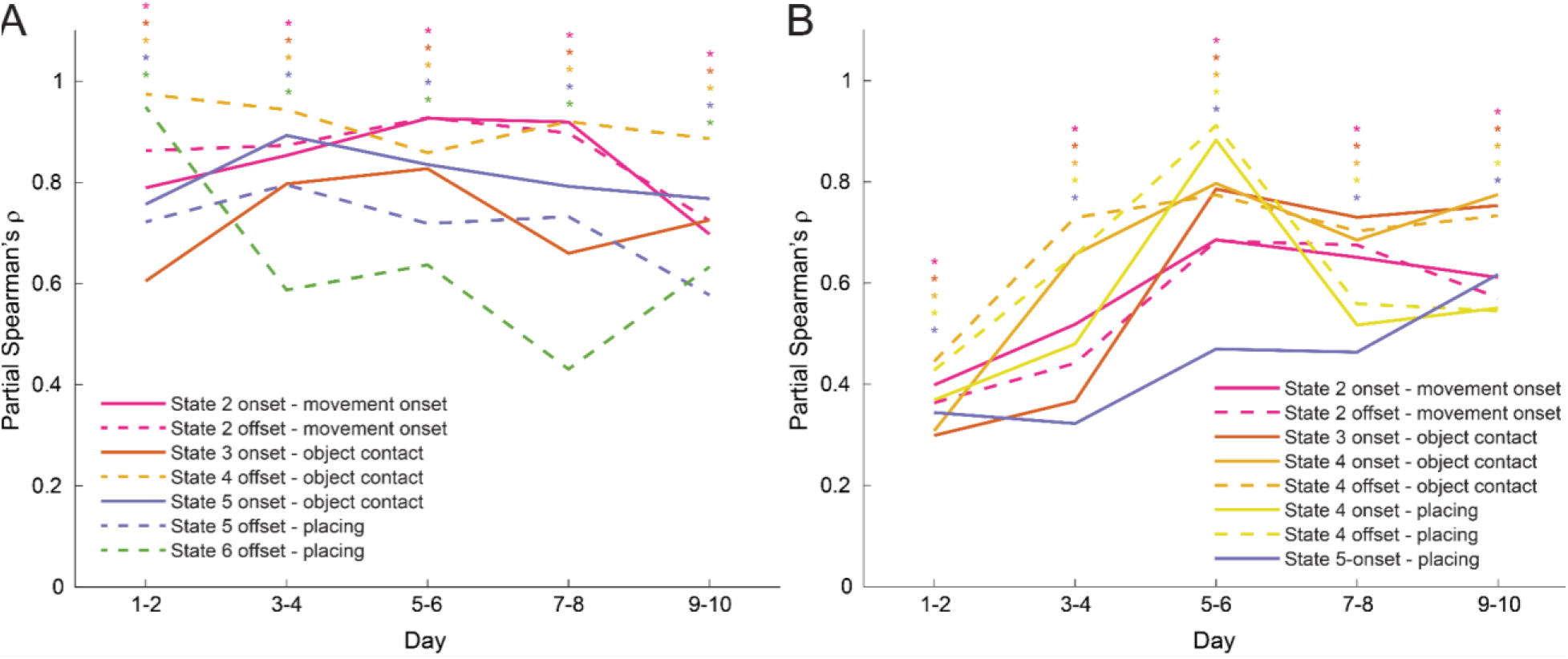
The link between states and behavioral events is consistent over days. **A)** Partial correlation coefficients between state transitions and behavioral events over the duration of recording from monkey 1, in groups of two days. Asterisks indicate statistically significant correlations. **B)** As in (A) for monkey 2.

### 3.3 M1 population state dynamics are related to behavioral state durations

We next compared response time (the time between the go signal and the hand movement onset events), reach duration (the time between the hand movement onset and object contact events), placing duration (the time between the object contact and placing events), and state dynamics between task conditions. States were characterized by their number of activations per trial and their lifetime (the time that state remains active, taken for each trial as the maximum duration over all activations in that trial).

There was no difference in response time between conditions (monkey 1: *X*^2^(2) = 0.66, *p* = 0.717; monkey 2: *X*^2^(2) = 4.37, *p* = 0.113), but conditions varied in terms of reach duration (monkey 1: *X*^2^(2) = 78.68, *p* < 0.001; monkey 2: *X*^2^(2) = 9.09, *p* = 0.011; Figure S5). For monkey 1, reach durations to the left target location (*M* = 408, *SE* = 8.40 ms) were longer than to the right (*M* = 343, *SE* = 8.58 ms, *t*(430) = 5.83, *p* < 0.001) and center (*M* = 312, *SE* = 8.48 ms, *t*(432) = −8.67, *p* < 0.001), and longer to the right compared to the center target location (*t*(436) = −2.79, *p* = 0.015). For monkey 2, reach durations to the left target (*M* = 230, *SE* = 13.2 ms) were shorter than to the right (*M* = 264, *SE* = 13.3, *t*(372) = −3.01, *p* = 0.008). Placing duration was only different for monkey 1 (*X*^2^(2) = 30.18, *p* < 0.001), and longer in the left (*M* = 641, *SE* = 15.7 ms) compared to the right (*M* = 582, *SE* = 15.8 ms, *t*(436) = 3.41, *p* = 0.002) and center (*M* = 548, *SE* = 15.8 ms, *t*(430) = −5.43, *p* < 0.001) conditions, but there was no difference between placing duration for the center and right conditions (*t*(433) = −1.96, *p* = 0.123).

The number of state activations per trial and their lifetime varied by state and condition in monkey 1 (activations: *X*^2^(10) = 95.94, *p* < 0.001; Figure 5; lifetime: *X*^2^(10) = 66.00, *p* < 0.001; Figure 6). State 1 had a shorter lifetime in the center (*M* = 533.5, *SE* = 11.4 ms) compared to the left (*M* = 635.5, *SE* = 11.2 ms, *t*(2616) = −6.39, *p* < 0.001) and right (*M* = 626.9, *SE* = 11.2 ms, *t*(2616) = −5.84, *p* < 0.001) conditions. State 3 was activated more times the left condition (*M* = 0.67, *SE* = 0.06) than in the center (*M* = 0.36, *SE* = 0.07, *z* = −3.37, *p* = 0.002) and right (*M* = 0.39, *SE* = 0.07, *z* = 3.12, *p* = 0.005) conditions. State 4 had longer duration in the left condition (*M* = 160.2, *SE* = 11.2 ms) than in the center (*M* = 97.7, *SE* = 11.4 ms, *t*(2616) = −3.92, *p* < 0.001) and right (*M* = 114.4, *SE* = 11.2, *t*(2616) = 2.89, *p* = 0.011) conditions. In contrast, state 5 was activated fewer times in the left condition (*M* = 0.11, *SE* = 0.08) than the center (*M* = 0.61, *SE* = 0.06, *z* = 5.10, *p* < 0.001) and right (*M* = 0.68, *SE* = 0.06, *z* = −5.81, *p* < 0.001) conditions. State 6 was also activated more times and had longer lifetime in the left condition (activations: M = 0.77, SE = 0.06; lifetime: M = 114.8, SE = 11.2 ms) than the center (activations: M = 0.26, SE = 0.07, z = −5.59, p < 0.001, lifetime: M = 43.1, SE = 11.4 ms, t(2616) = −4.50, p < 0.001) and right (activations: M = 0.30, SE = 0.07, z = 5.23, p < 0.001; lifetime: M = 67.7, SE = 11.2 ms, t(2616) = 2.97, p = 0.008) conditions. The dynamics of state 2 did not vary by condition. In monkey 2, there was only a main effect of state on the number of state activations per trial and their lifetime, however, there was almost no difference between conditions in response time, reach duration, and placing duration for this monkey.

**Figure 5.**
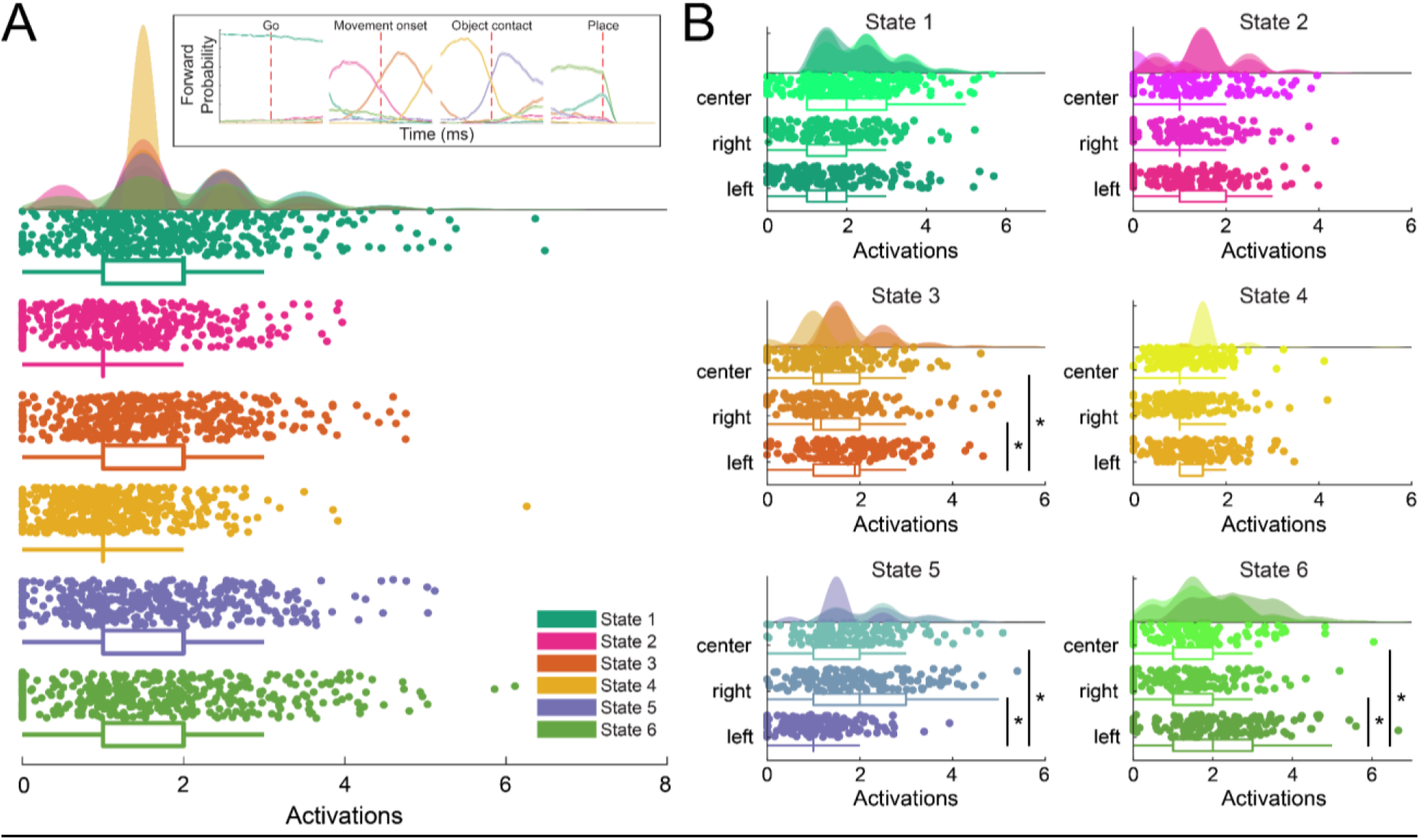
Number of state activations per trial compared between conditions. **A)** The number of activations per trial for each state identified by the model. Shaded areas indicate the distribution density, scatter plots show the number of activations for individual trials, and box plots show descriptive statistics: median (in the box), interquartile interval (the box), outliers (above whiskers). The inset shows the event-aligned mean state forward probabilities from Figure 2. **B)** Number of activations per trial for each state compared by conditions: center, right, left. Asterisks indicate significant pairwise comparisons.

**Figure 6.**
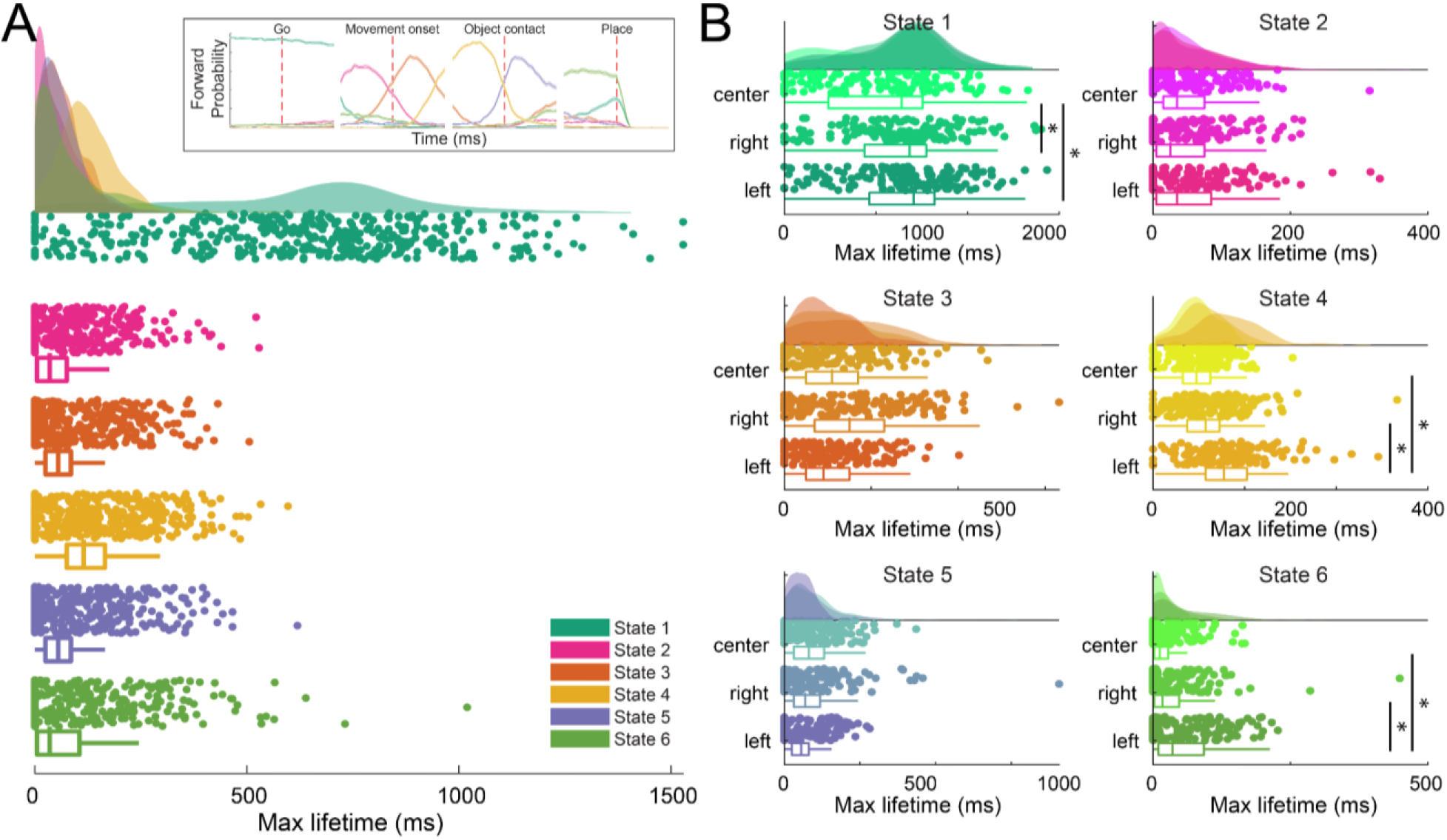
Maximum state lifetime compared between conditions. **A)** The maximum lifetime (duration of the longest activation per trial) for each state identified by the model. Shaded areas indicate the distribution density, scatter plots show the maximum state lifetime for individual trials, and box plots show descriptive statistics: median (in the box), interquartile interval (the box), outliers (above whiskers). The inset shows the event-aligned mean state forward probabilities from Figure 2. **B)** Maximum lifetime for each state compared by conditions: center, right, left. Asterisks indicate significant pairwise comparisons.

Finally, we looked at relationships between state dynamics and response time, reach duration, and placing duration. For monkey 1, response time was correlated with the number of activations (*ρ* = 0.11, *p* = 0.03) and lifetime (*ρ* = 0.46, *p* = 0.005) of state 1. Reach duration was correlated with the number of activations of states 3 (*ρ* = 0.15, *p* = 0.002), 4 (*ρ* = 0.23, *p* < 0.001), and 6 (*ρ* = 0.17, *p* = 0.001). Placing duration was correlated with the number of activations of states 1 (*ρ* = 0.17, *p* = 0.001) and 4 (activations: ρ = 0.11, p = 0.018), as well as the number of activations and lifetime of state 6 (activations: *ρ* = 0.34, *p* < 0.001; lifetime: *ρ* = 0.39, *p* = 0.020). For monkey 2, response time was correlated with the number of activations (*ρ* = 0.34, *p* < 0.001) and lifetime (*ρ* = 0.22, *p* = 0.005) of state 1, and the number of activations of state 2 (*ρ* = 0.17, *p* < 0.001). Reach duration was correlated with the number of activations of states 2 (*ρ* = 0.23, *p* < 0.001), 3 (*ρ* = 0.25, *p* < 0.001), and 5 (*ρ* = 0.22, *p* < 0.001), and the number of activations and lifetime of state 4 (activations: *ρ* = 0.20, *p* < 0.001; lifetime: *ρ* = 0.22, *p* = 0.003). Placing duration was correlated with the number of activations of state 2 (*ρ* = 0.22, *p* < 0.001), the lifetime of state 4 (*ρ* = 0.22, *p* = 0.003), and the number of activations and lifetimes of states 3 (activations: *ρ* = 0.30, *p* < 0.001; lifetime: *ρ* = 0.15, *p* = 0.036), and 5 (activations: *ρ* = 0.37, *p* < 0.001; lifetime: *ρ* = 0.18, *p* = 0.015). The number of activations and lifetime of identified states was therefore related to response time, reach duration, and placing duration in a state-specific way, with earlier occurring states related to response time, states occurring in the middle of the trial to reach duration, and states occurring later to placing duration.

### 3.4 States represent distinct spatiotemporal patterns of neural population activity

Having determined that the timing, number of activations, and lifetime of the HMM-identified states are tightly coupled to the behavioral phases of reaching, grasping, and placing an object, we next examined neural dynamics associated with state transitions and occurring within states. We aligned multi-unit firing rates to the onset and offset times of each state in every trial, and baseline-corrected them using activity from the 100 ms prior to the state onset (monkey 1: Figure 7; monkey 2: Figure S6). State transitions tended to be associated by rapid changes in firing rates, followed by state-specific patterns of activity. For each electrode, we compared the effect of state on the multi-unit firing rate in the 100 ms prior to state onset to that in the first half of the state duration, and the multi-unit firing rate in the last half of the state duration with that in the 100 ms following the state offset. For monkey 1, all electrodes had a significant interaction between state and time period, and follow-up comparisons showed that states differed in the number of electrodes in which multi-unit firing changed with the onset (state 1: *N* = 0, state 2: *N* = 14, state 3: *N* = 18, state 4: *N* = 19, state 5: *N* = 19, state 6: *N* = 16) and offset (state 1: *N* = 16, state 2: *N* = 20, state 3: *N* = 15, state 4: *N* = 19, state 5: *N* = 14, state 6: *N* = 18) of the state. The same was true for monkey 2 for all electrodes for both state onset (state 1: *N* = 0, state 2: *N* = 18, state 3: *N* = 24, state 4: *N* = 15, state 5: *N* = 20) and offset (state 1: *N* = 20, state 2: *N* = 19, state 3: *N* = 21, state 4: *N* = 21, state 5: *N* = 12). In order to determine if activity within each state represented dynamic or sustained patterns of activity, we similarly compared the effect of state on firing rate in the first half of the state duration to that in the second half. A significant interaction between state and time period was found in 19 out the 21 electrodes for monkey 1, with the fewest number of electrodes exhibiting within-state dynamic activity in state 1 and the greatest number in state 4 (state 1: *N* = 0, state 2: *N* = 5, state 3: *N* = 9, state 4: *N* = 19, state 5: *N* = 9, state 6: *N* = 6). For monkey 2, there was a significant state by time period interaction in 24 out of the 25 electrodes, with the least within-state dynamics in state 1 and the most in state 5 (state 1: *N* = 5, state 2: *N* = 15, state 3: *N* = 14, state 4: *N* = 16, state 5: *N* = 14). In both monkeys, state 1 therefore represents a static period of low activity in all electrodes, but each other state represents a transition to a specific combination of sustained and dynamic activity across electrodes.

**Figure 7.**
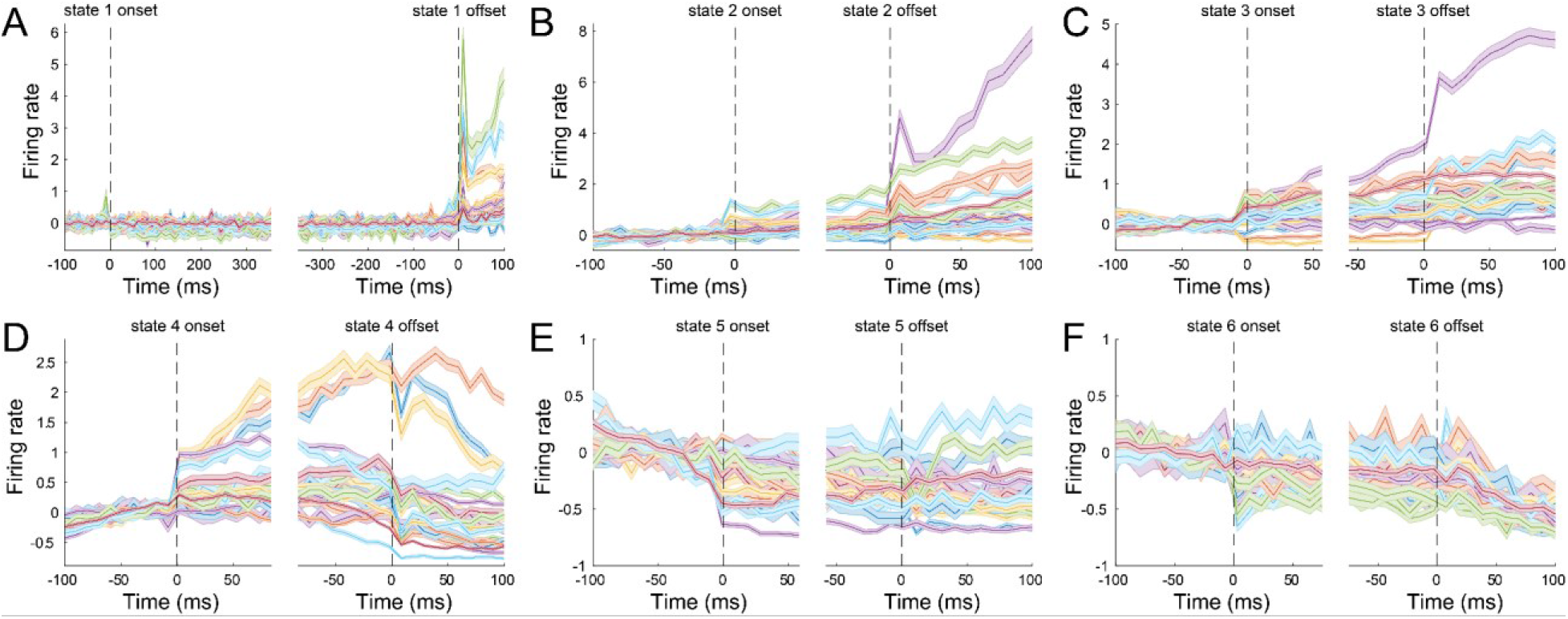
States represent different patterns of neural activity patterns. Each panel shows mean, baseline-corrected firing rates from each electrode from monkey 1 aligned to the onset (left plot) and offset (right plot) of states 1-6 (A-F). Dashed vertical lines indicate the onset and offset time of the state. State transitions are associated with rapid changes in state-specific patterns of neural activity.

We then evaluated the spatiotemporal specificity of these activity patterns by shuffling the data, either temporally (i.e. shuffling time points; Figure S7) or spatially (i.e. shuffling electrodes; Figure S8) and using the HMM to obtain forward probabilities and sequences of the most likely state. For monkey 1, both temporal and spatial shuffling abolished the relationships between the movement onset time and the onset (temporal shuffled *p* = 0.14; compared to unshuffled: *ρ* = 0.01; spatial shuffled *ρ* = 0.08; compared to unshuffled: *p* = 0.01) and offset (temporal shuffled ρ = 0.09; compared to unshuffled: p = 0.01; spatial shuffled ρ = 0.04; compared to unshuffled: p = 0.01) of state 2. The correlations between the object contact event and the onset of state 3 (temporal shuffled *ρ* = −0.02, compared to unshuffled: *p* = 0.02; spatial shuffled *ρ* = 0.10, compared to unshuffled: *p* = 0.02) and the onset of state 5 (temporal shuffled *ρ* = 0.10, compared to unshuffled: *p* = 0.01; spatial shuffled *ρ* = 0.13, compared to unshuffled: *p* = 0.01) were also destroyed by temporal and spatial shuffling. Only spatial shuffling had an effect on the correlation between the placing event and the onset of state 6 (spatial shuffled *ρ* = −0.08, compared to unshuffled: *p* = 0.02), and only temporal shuffling disrupted the relationship between object contact and the offset of state 4 (*ρ* = −0.02, *p* = 0.01). For monkey 2, both temporal and spatial shuffling abolished the relationships between the movement onset time and the onset (temporal shuffled *ρ* = −0.05; compared to unshuffled: *p* = 0.01; spatial shuffled *ρ* = 0.06; compared to unshuffled: *p* = 0.01) and offset (temporal shuffled ρ = −0.05; compared to unshuffled: *p* = 0.01; spatial shuffled ρ = 0.04; compared to unshuffled: p = 0.01) of state 2. The correlations between the object contact event and the onset of state 3 were not destroyed by either temporal or spatial shuffling, but those between object contact and the onset (temporal shuffled *ρ* = −0.03, compared to unshuffled: *p* = 0.01; spatial shuffled *ρ* = 0.10, compared to unshuffled: *p* = 0.02) and offset (temporal shuffled *ρ* = −0.02, compared to unshuffled: *p* = 0.01; spatial shuffled *ρ* = 0.06, compared to unshuffled: *p* = 0.01) of state 4 were affected by both forms of shuffling. Both temporal and spatial shuffling had an effect on the correlation between the placing event and the offset of state 4 (temporal shuffled *ρ* = −0.03, compared to unshuffled: *p* = 0.01; spatial shuffled *ρ* = 0.04, compared to unshuffled: *p* = 0.01), but neither had an effect on that between the placing event and the onset of state 5. The states identified by the HMM therefore represent rapid transitions between distinct spatiotemporal patterns of neural population activity which are generated during specific phases of the reach, grasp, and place task.

## 4. Discussion

In this paper we present a new multilevel Bayesian HMM, designed for analyzing longitudinal neural recordings from chronic multi-electrode arrays through the use of trial-specific random effects in transition and emission probabilities, multivariate Poisson log-normal emission probability distributions, and trial-specific condition covariates. We trained instantiations of the model on long term data recorded from the primary motor cortex of the macaque monkey during a task involving reaching, grasping, and placing a target object. We showed that although the model was not trained with any information about the task structure, the timing of discrete state transitions was tightly linked to behavioral events (i.e. the timing of the movement onset, object contact, and placing events), and state dynamics were closely aligned with the duration of intervals between events (i.e. response time, reach duration, placing duration). The states identified by the HMM represented fast transitions between distinct spatiotemporal patterns of neural population activity which are generated during specific phases of the reach, grasp, and place task, likely corresponding to different motor control processes.

At the single trial level, various states were coupled to different phases of the task. In both monkeys, state 1 represented a static period of low activity across all electrodes, and the go signal event did not elicit any state transition. However, sometime after the go signal, there was a transition to state 2, whose onset and offset correlated with movement onset time. State 2 therefore likely represents movement preparation or initiation mechanisms. States 3, 4, and 5 in monkey 1 and states 3 and 4 in monkey 2 were correlated with the object contact, the reach duration, and the placing duration and therefore may be differentially involved in the acceleration and deceleration phases of reaching (Kadmon Harpaz et al., 2019), or the coordination of the wrist orientation. State 6 is activated before, and aligned to, the placing event, and therefore likely reflects control of a reaching movement in the opposite direction from the initial reach-to-grasp (Kadmon Harpaz et al., 2019). The model was trained solely on the multi-unit spike data from each trial without any event timing information, and yet state transitions and lifetimes were tightly linked to kinematic events and movement durations, suggesting that each state represents distinct motor control mechanisms.

There are a few limitations to this framework which include the basic assumptions of HMMs, longitudinal generalizability for predictive decoding, task conditions, zero-inflation, and the use of multi-unit, rather than single-unit, activity. All HMMs make the assumption that only one state can be active at any given time, and that state transition probabilities only depend on the current state rather than the recent history of states (the Markov assumption). These may not be reasonable assumptions for neural population activity, and there have been recent efforts to develop a similar framework without, or with relaxed versions of, these assumptions (Gohil et al., 2022). A promising future direction might be to extend the benefits of multilevel Bayesian HMMs to such frameworks, for example by implementing an explicit-duration hidden semi-Markov model that relaxes the Markov assumption, decoupling the state duration from the transition probabilities by explicitly assigning distributions to the duration of the states (Yu, 2010). One advantage of the multilevel approach is the ability to model between-trial differences in activity that might undermine attempts to identify the population state. However, as would also be the case for single level models, new trials would have to use group-level mean model parameters. One possibility for future work might be to take advantage of the trial-specific parameters in the multilevel model and extrapolate them for decoding future activity. The task used involves only three reach directions and one grasp type. This was a necessary restriction for the current study to establish and test the framework, but future applications should use a wider range of reach and grasp movements along with kinematic tracking in order to identify the specific motor control mechanisms represented by each state. Model checking with PPCs revealed a minor underestimation of the proportion of zeros in the spike count data, consistent with a zero-inflated generative process. Future versions of the model could be extended with a zero-inflated Poisson emission distribution to model the excess of zeros beyond what a common random-effect Poisson distribution can accommodate (Aguero-Valverde, 2013). Finally, we trained the model on multi-unit spiking data, due to the inherent difficulty in tracking single units over long-term recordings. However, this is also a strength of this approach, as multi-unit activity has been shown to be sufficient to capture neural population dynamics (Trautmann et al., 2019), and therefore circumvents the need for longitudinal spike sorting.

## 5. Conclusion

The multilevel Bayesian HMM presented here has several unique advantages for analyzing longitudinal neural population dynamics from multi-electrode arrays compared to current approaches. The emission probabilities are multivariate Poisson distributions, capturing the statistical distribution of neuronal spikes. The model is trained trial-by-trial, without the need for epochs of the same length, making it applicable to motor tasks which typically have variable trial duration due to movement variability. The inclusion of covariates allows the model to be fit on data from multiple experimental conditions and therefore simplifies comparison between them. In this study, each monkey was pre-trained on the reaching, grasping, and placing task, but because the multilevel Bayesian framework easily handles longitudinal data, it is especially well suited for studies of longer term neural population plasticity in the context of motor learning.

## Acknowledgements

We acknowledge funding from the European Research Council (ERC) under the European Union’s Horizon 2020 research and innovation programme (ERC-CoG 864550 to JB), and the National Institutes of Health (NICHD P01HD064653 to PFF). This work made use of the Dutch national e-infrastructure Snellius with the support of the SURF Cooperative using grant no. EINF-2570, which is (partly) financed by the Dutch Research Council (NWO). SK was supported by the Fondation pour la Recherche Médical (FRM). We thank H Scherberger and PA Beuriat for assistance with array implantation. We thank M Montagna and S Pinède for their help with the construction of the laboratory set up.

## Supplementary material

**Figure S1.**
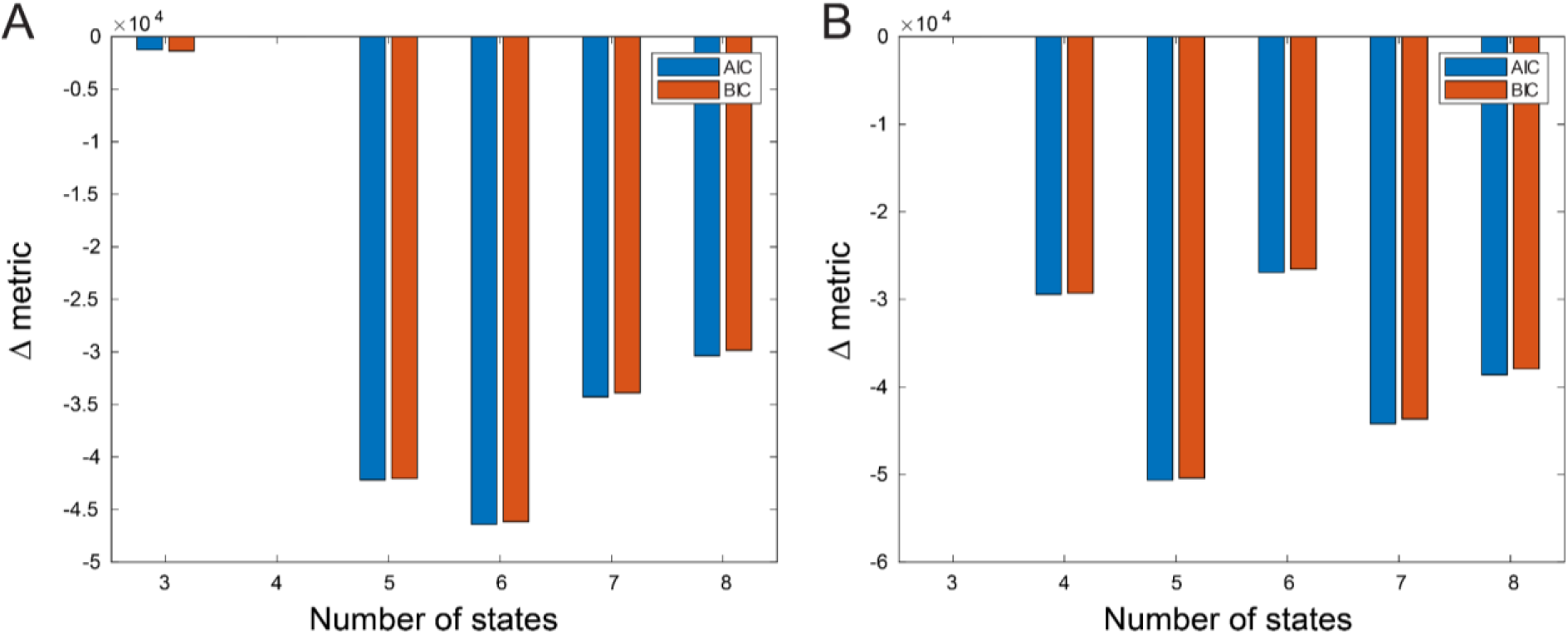
Model comparison results. AIC and BIC, relative to the worst model, for monkey 1 (A) and monkey 2 (B) for models with 3 to 8 states. For monkey 1, the best fitting model had 6 states, whilst a 5 state model had the best fit for monkey 2.

**Figure S2.**
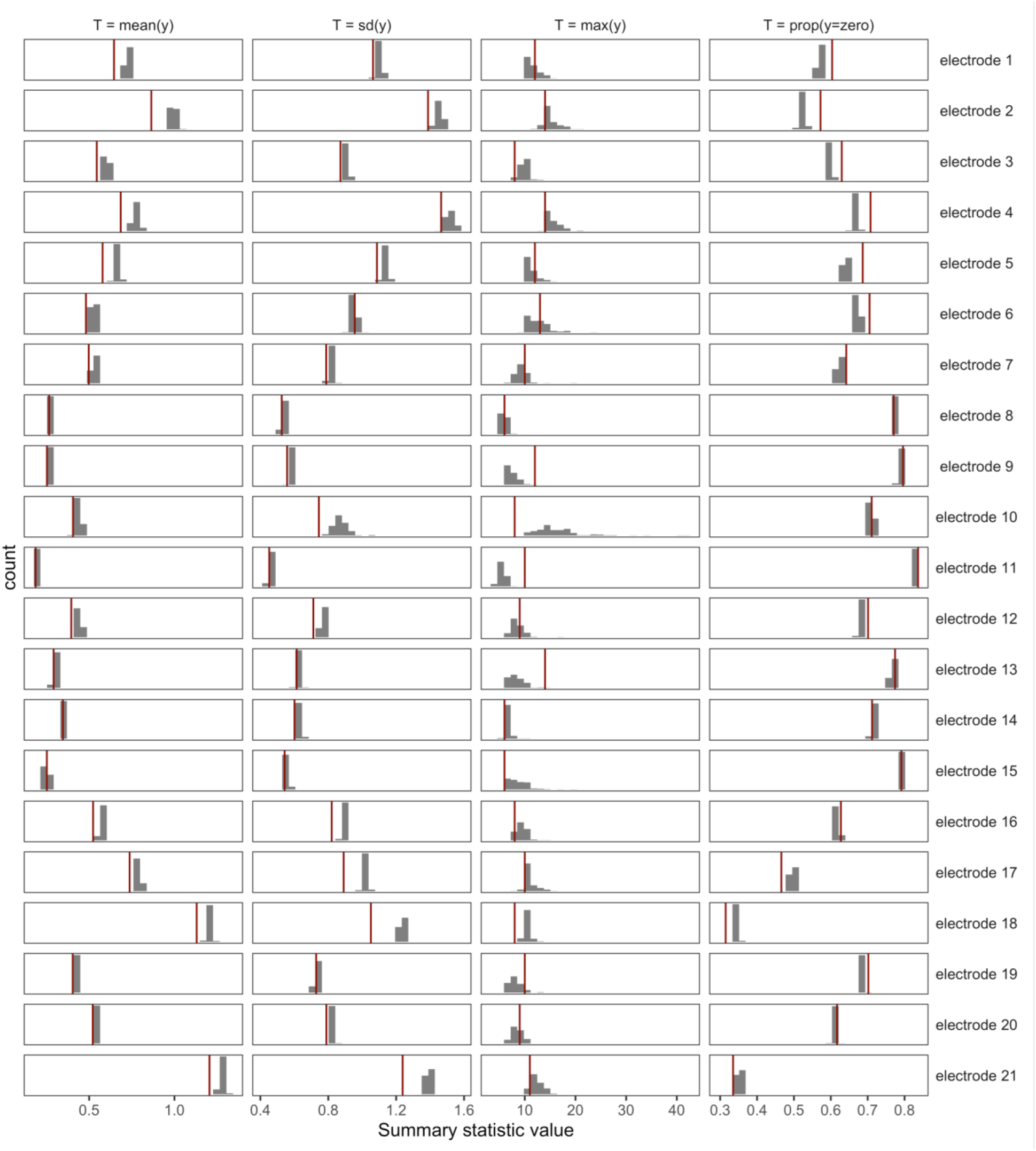
The multilevel Bayesian HMM is able to reproduce the general patterns of four summary statistics on the aggregated empirical data of the first monkey. Histograms show the distribution of four summary statistics *T(y)_rep_* (mean, standard deviation, max, and proportion of zeros) of the synthetic spike counts for the 21 electrodes from monkey 1 over 500 simulated data sets. The red vertical lines indicate the actual scores of the summary statistics *T(y)_emp_* from the empirical data. The synthetic data simulated by the multilevel HMM was able to recover the general pattern of summary statistics of the aggregated empirical data set, indicating adequate fit to the data.

**Figure S3.**
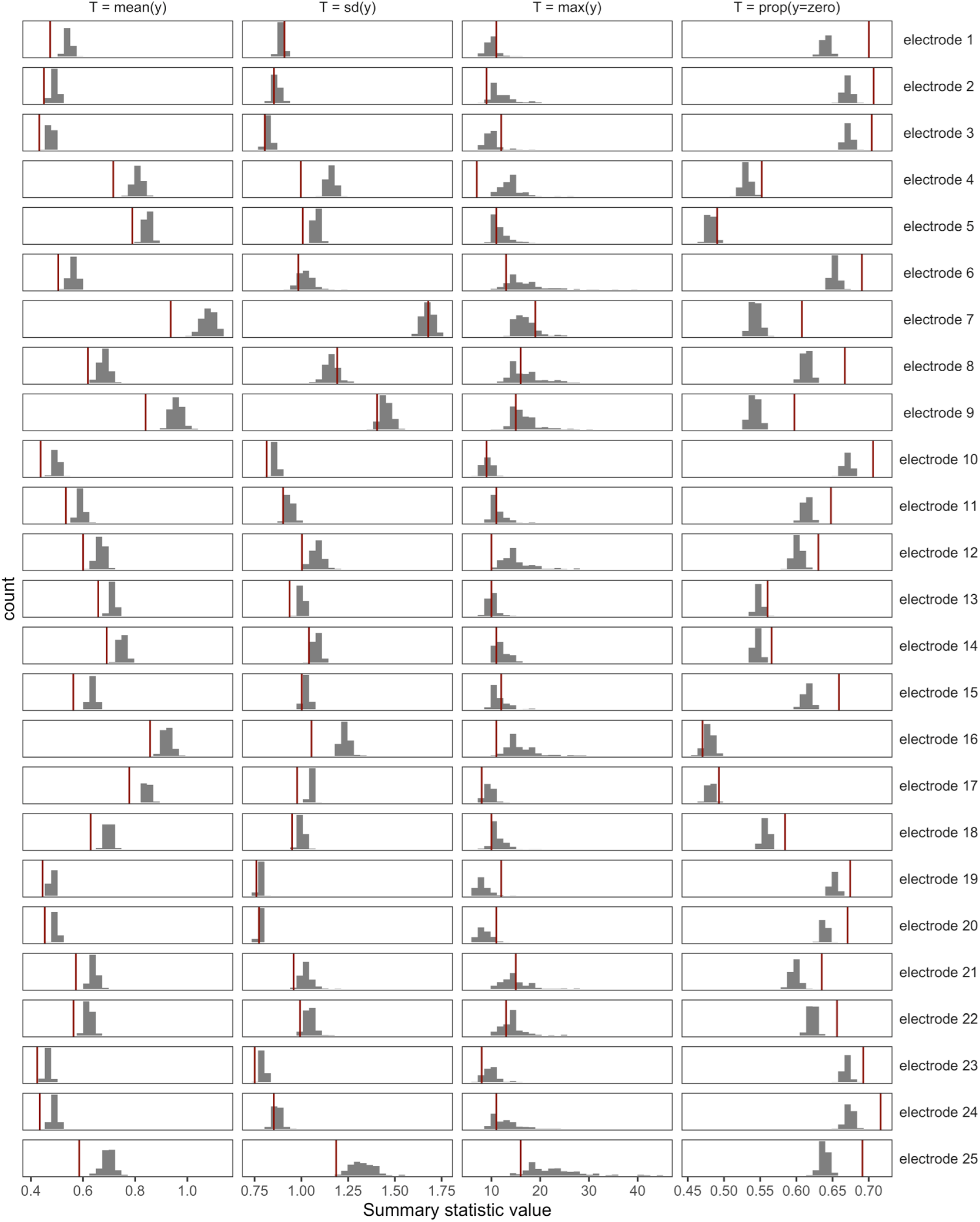
The multilevel Bayesian HMM is able to reproduce the general patterns of four summary statistics on the aggregated empirical data of the second monkey. Histograms show the distribution of four summary statistics *T(y)_rep_* (mean, standard deviation, max, and proportion of zeros) of the synthetic spike counts for the 25 electrodes from monkey 2 over 500 simulated data sets. The red vertical lines indicate the actual scores of the summary statistics *T(y)_emp_* from the empirical data. The synthetic data simulated by the multilevel HMM was able to recover the general pattern of summary statistics of the aggregated empirical data set, indicating adequate fit to the data.

**Figure S4.**
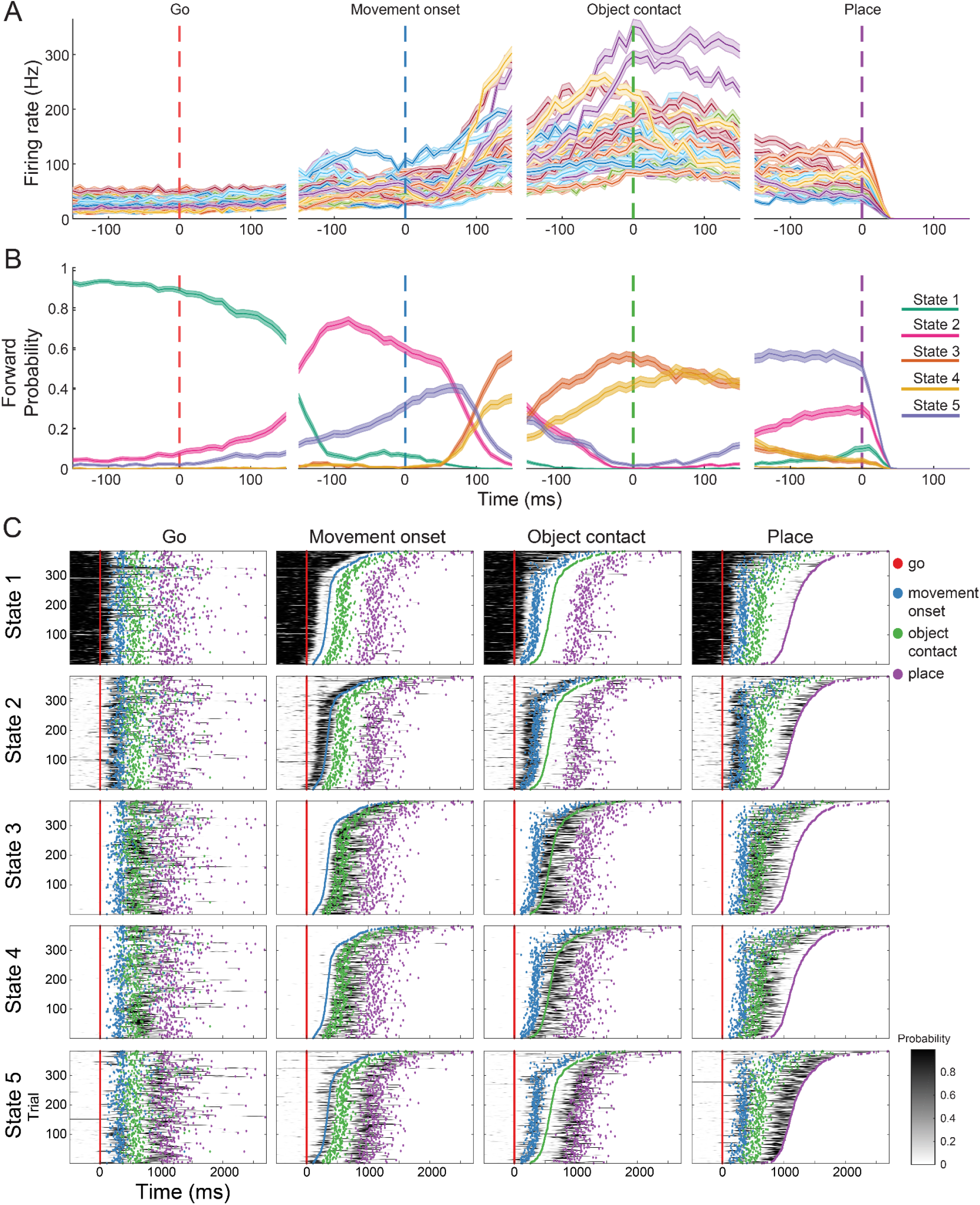
States identified from monkey 2 are also tightly linked to behavioral events. **A)** Mean multi-unit firing rate of 25 electrodes from the M1 array of monkey 2 over 10 days (383 trials) aligned to behavioral events (go: go signal tone, hand movement onset: beginning of the reaching movement, object contact: grasping of the object, place: placing the object). **B)** Mean forward probabilities generated by the model for each state, averaged over the same trials and aligned to the same behavioral events as in (A). **B)** Each row shows the forward probabilities from one of the model states for each trial, over all days of recording, for monkey 2. Each column shows the trial forward probabilities sorted by one of the behavioral events (from left to right: go signal, hand movement onset, object contact, and placing).

**Figure S5.**
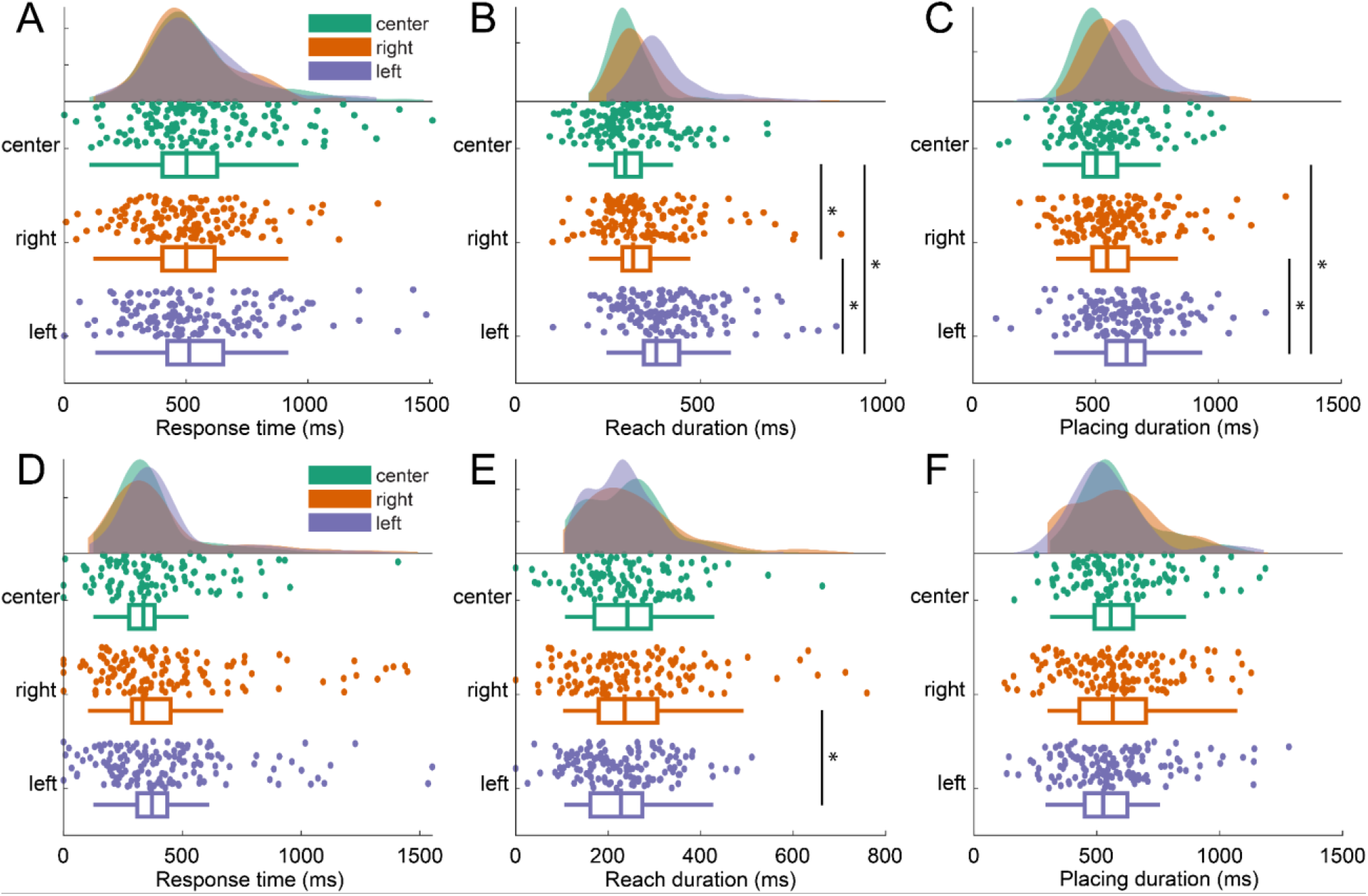
Response time, reach and place durations, but not response time vary by condition. The response time (A), reach duration (B), and placing duration (C) for each trial from monkey 1, and monkey 2 (D-F), by condition. Shaded areas indicate the distribution density, scatter plots show the values for individual trials, and box plots show descriptive statistics: median (in the box), interquartile interval (the box), outliers (above whiskers). Asterisks indicate significant pairwise comparisons.

**Figure S6.**
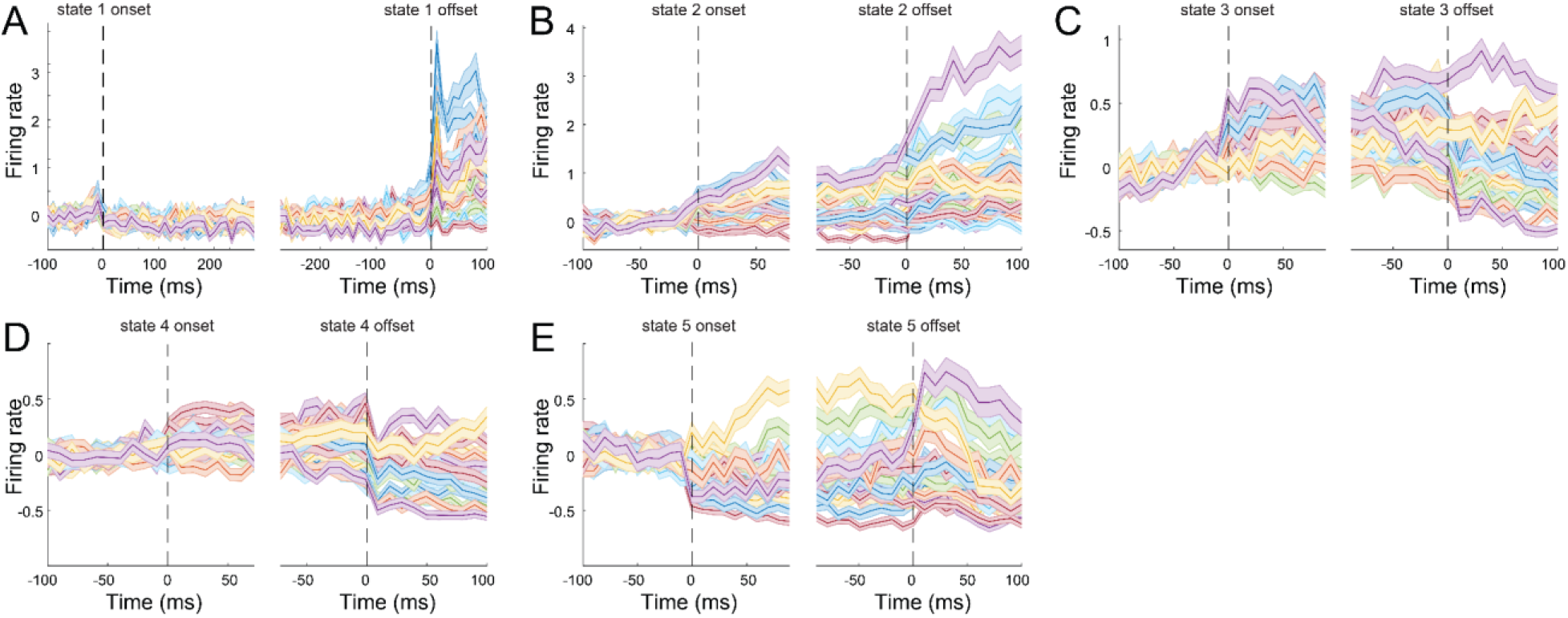
States represent different patterns of neural activity patterns from monkey 2. Each panel shows mean, baseline-corrected firing rates from each electrode from monkey 2 aligned to the onset (left plot) and offset (right plot) of states 1-6 (A-F). Dashed vertical lines indicate the onset and offset time of the state. State transitions are associated with rapid changes in state-specific patterns of neural activity.

**Figure S7.**
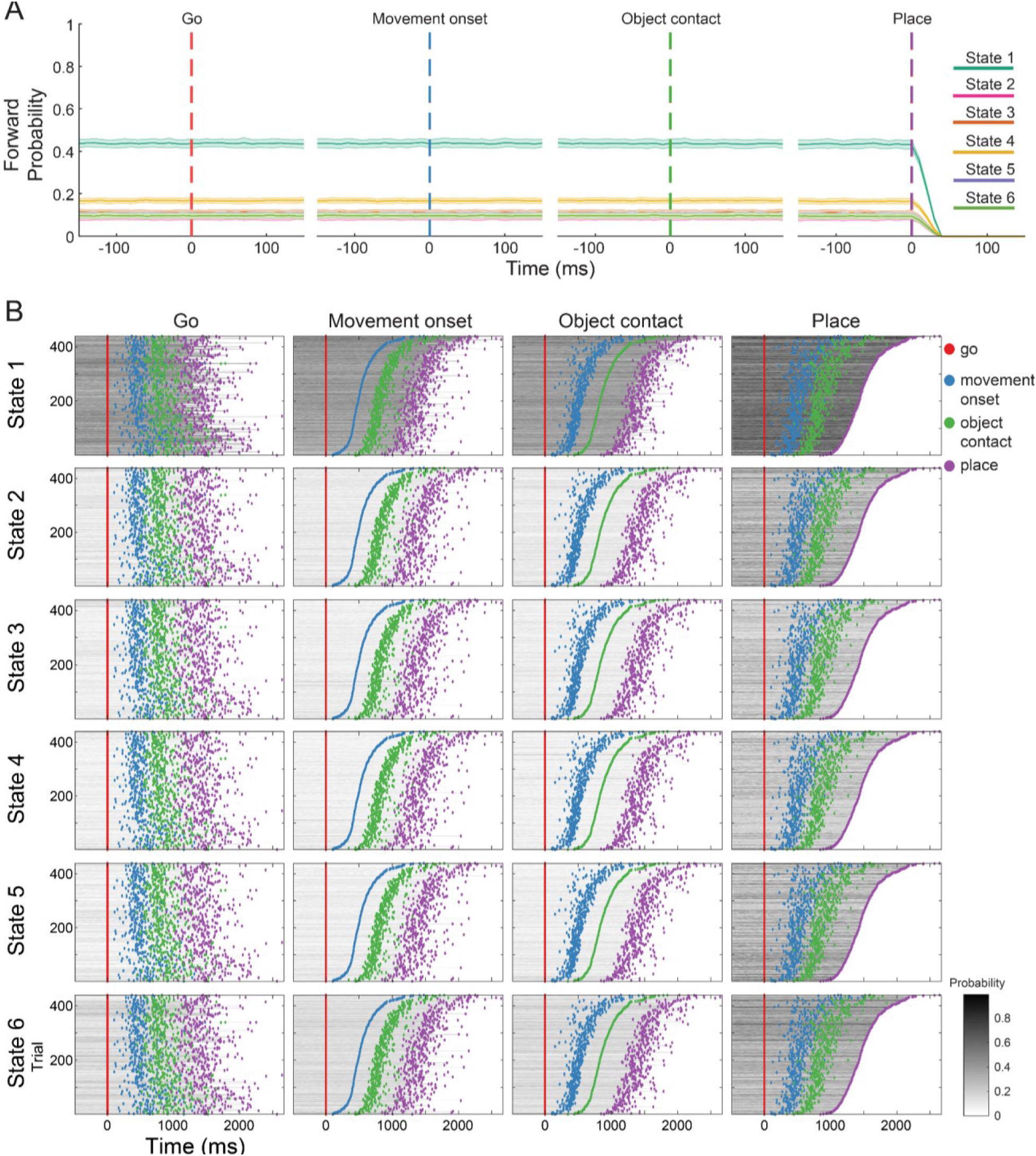
Temporal shuffling destroys the coupling between states and behavioral events. **A)** Mean forward probabilities generated by the model for monkey 1 for each state, averaged over 100 iterations of temporal shuffling and aligned to the behavioral events. **B)** Each row shows the forward probabilities from one of the model states for each trial, averaged over 100 iterations of temporal shuffling. Each column shows the trial forward probabilities sorted by one of the behavioral events (from left to right: go signal, hand movement onset, object contact, and placing).

**Figure S8.**
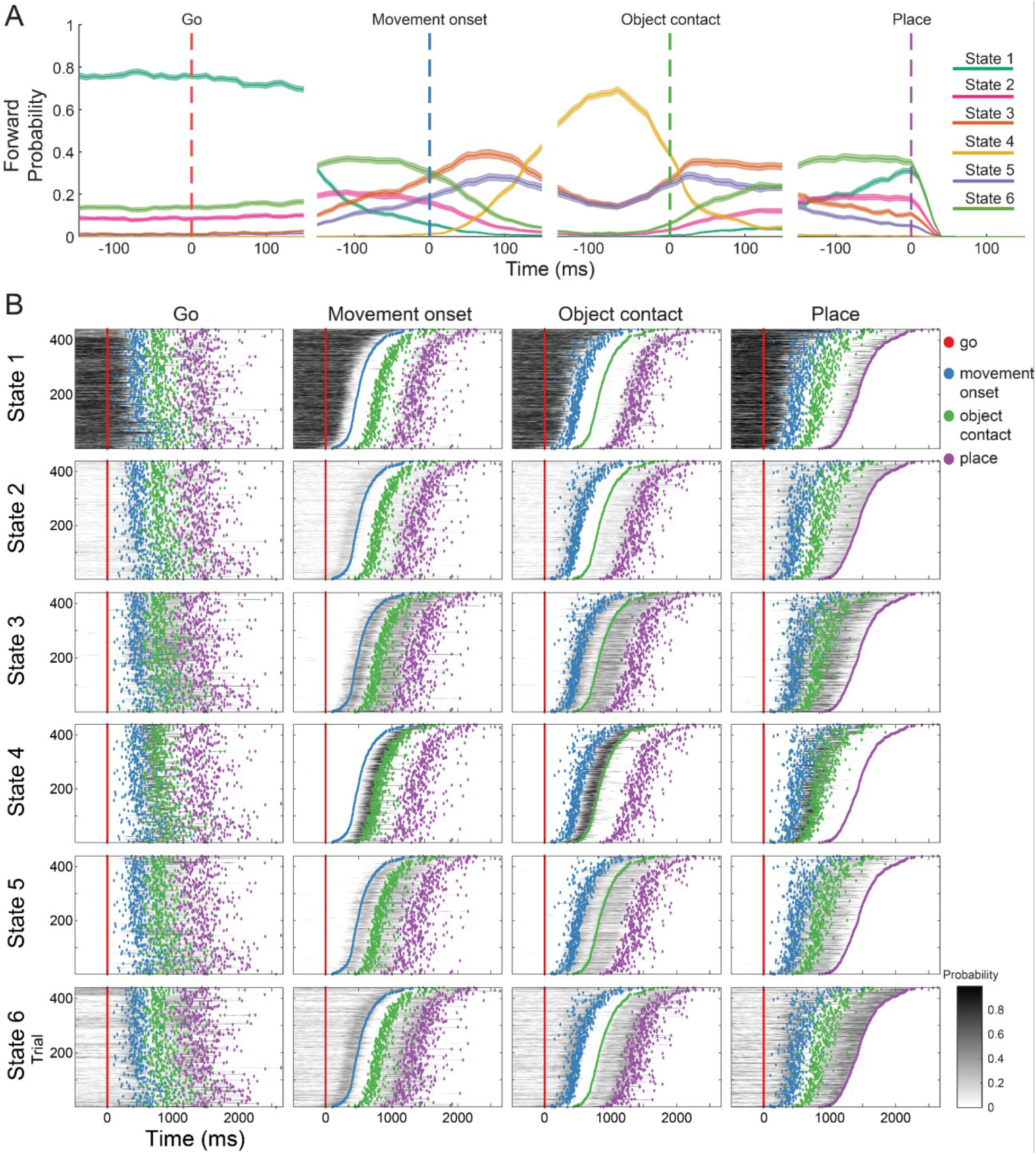
Spatial shuffling destroys the coupling between states and behavioral events. **A)** Mean forward probabilities generated by the model for monkey 1 for each state, averaged over 100 iterations of spatial (electrode) shuffling and aligned to the behavioral events. **B)** Each row shows the forward probabilities from one of the model states for each trial, averaged over 100 iterations of spatial (electrode) shuffling. Each column shows the trial forward probabilities sorted by one of the behavioral events (from left to right: go signal, hand movement onset, object contact, and placing).

**Table S1.** Mean evaluation metrics for the results of the Monte Carlo simulation study on the state decoding accuracy for each monkey.

| Metric | Mean | SD | CI2.5% | CI97.5% |
| --- | --- | --- | --- | --- |
| **Monkey 1** |  |  |  |  |
| Accuracy | 0.903 | 0.002 | 0.899 | 0.908 |
| Bal. Accuracy | 0.916 | 0.002 | 0.913 | 0.920 |
| F1 score | 0.846 | 0.004 | 0.839 | 0.853 |
| Cohen's Kappa | 0.863 | 0.003 | 0.857 | 0.869 |
| **Monkey 2** |  |  |  |  |
| Accuracy | 0.847 | 0.009 | 0.829 | 0.864 |
| Bal. Accuracy | 0.875 | 0.006 | 0.863 | 0.887 |
| F1 score | 0.781 | 0.010 | 0.762 | 0.800 |
| Cohen's Kappa | 0.784 | 0.011 | 0.762 | 0.804 |

**Table S2.**
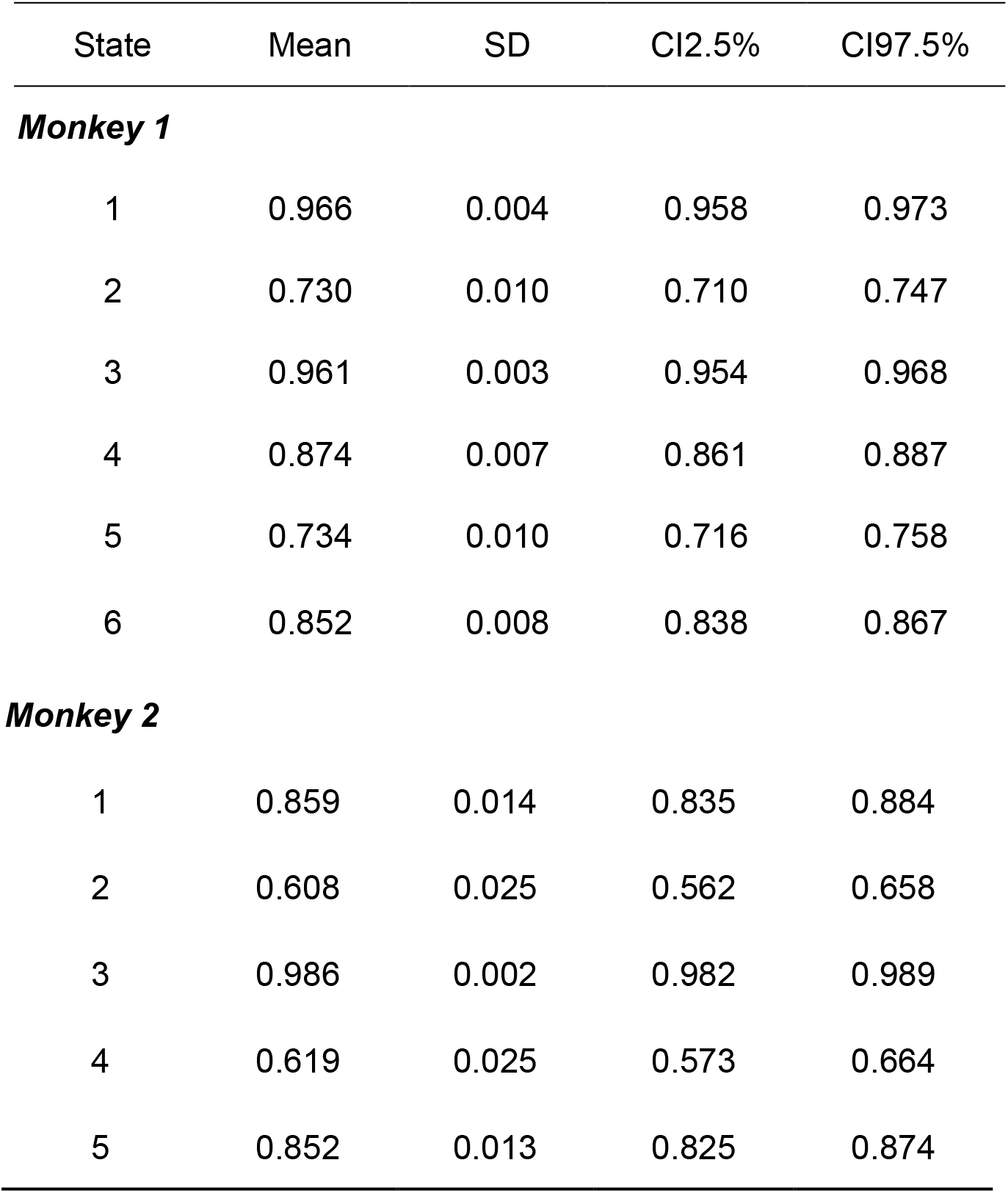
Mean decoding accuracy by state for the results of the Monte Carlo simulation study for each monkey.

| State | Mean | SD | CI2.5% | CI97.5% |
| --- | --- | --- | --- | --- |
| **Monkey 1** |  |  |  |  |
| 1 | 0.966 | 0.004 | 0.958 | 0.973 |
| 2 | 0.730 | 0.010 | 0.710 | 0.747 |
| 3 | 0.961 | 0.003 | 0.954 | 0.968 |
| 4 | 0.874 | 0.007 | 0.861 | 0.887 |
| 5 | 0.734 | 0.010 | 0.716 | 0.758 |
| 6 | 0.852 | 0.008 | 0.838 | 0.867 |
| ***Monkey 2*** |  |  |  |  |
| 1 | 0.859 | 0.014 | 0.835 | 0.884 |
| 2 | 0.608 | 0.025 | 0.562 | 0.658 |
| 3 | 0.986 | 0.002 | 0.982 | 0.989 |
| 4 | 0.619 | 0.025 | 0.573 | 0.664 |
| 5 | 0.852 | 0.013 | 0.825 | 0.874 |

